# Tau accelerates tubulin exchange in the microtubule lattice

**DOI:** 10.1101/2024.10.05.616777

**Authors:** Subham Biswas, Rahul Grover, Cordula Reuther, Chetan S. Poojari, M. Reza Shaebani, Mona Grünewald, Amir Zablotsky, Jochen S. Hub, Stefan Diez, Karin John, Laura Schaedel

**Affiliations:** Experimental Physics and Center for Biophysics, Saarland University, 66123 Saarbrücken, Germany; B CUBE - Center for Molecular Bioengineering, TUD Dresden University of Technology, 01307 Dresden, Germany; Cluster of Excellence Physics of Life, TUD Dresden University of Technology, 01062 Dresden, Germany; Max Planck Institute of Molecular Cell Biology and Genetics, 01307 Dresden, Germany; Theoretical Physics and Center for Biophysics, Saarland University, 66123 Saarbrücken, Germany; Université Grenoble Alpes, CNRS, LIPhy, 38000 Grenoble, France

## Abstract

Microtubules are cytoskeletal filaments that exhibit dynamic tip instability and, as recent discoveries reveal, possess a dynamic lattice undergoing continuous tubulin loss and incorporation. In this study, we investigate the role of tau, a neuronal microtubule-associated protein (MAP) known for its stabilizing effects on microtubules, in modulating lattice dynamics. Using in vitro reconstitution, kinetic Monte Carlo modeling, and molecular dynamics simulations, we reveal that tau, despite lacking enzymatic activity, accelerates tubulin exchange within the lattice, particularly at topological defect sites. Tau appears to stabilize longitudinal tubulin–tubulin interactions while destabilizing lateral ones, thereby enhancing the mobility and repair of lattice defects. These results challenge the traditional view of tau as merely a stabilizer, uncovering its active role in dynamically modulating microtubule lattice structure.

## Introduction

Microtubules are an essential filamentous component of the cytoskeleton. They exhibit unique dynamic and mechanical properties owing to their hollow structure, their highly ordered arrangement of tubulin dimer subunits, and their dissipative growth dynamics. Traditionally, research has focused on the dynamic behavior of microtubule tips, whose intrinsic instability facilitates rapid growth and shrinkage^1^. In contrast, the microtubule shaft (often termed ‘lattice’), was long considered a static structure and thus received less attention in studies on the dynamic regulation of microtubules.

Recent discoveries have challenged this view, revealing that the microtubule lattice is far from static. Tubulin dimers continuously incorporate into and dissociate from the lattice, rendering it dynamic^2^. Importantly, this dynamicity endows microtubules with the capacity for self-repair and resilience against mechanical stresses typical in cellular environments, such as bending^3^ and friction at crossing points^4,5^. The dynamic turnover within the lattice in the absence of mechanical forces has been attributed to defect sites^6–8^, which act as focal points for tubulin exchange due to the missing inter-subunit bonds of the surrounding tubulin dimers^2,9^. Although lattice dynamics can occur independently of other cellular factors and are thus intrinsic to microtubules, they can be modulated by microtubule-associated proteins (MAPs) such as CLIP-170^5^, CLASP^10^, molecular motors^11–13^, and severing enzymes^14^.

Tau, a well-studied neuronal MAP, predominantly decorates axonal microtubules^15–17^ and is known for its role in microtubule nucleation, bundling, and stabilization against depolymerization^18–23^. Although tau’s influence on microtubule tip dynamics and microtubule bundling has been extensively studied, its impact on lattice dynamics has not been addressed so far. Tau binds along the length of microtubules, interacting with multiple tubulin dimers^24^. This suggests that it could significantly affect the stability of the lattice.

In this study, we explore the role of tau in microtubule lattice dynamics. Through a combination of in vitro reconstitution experiments and kinetic Monte Carlo modeling, supported by molecular dynamics simulations, we discover that tau, while generally stabilizing microtubules, surprisingly accelerates the exchange of tubulin in the microtubule lattice. This exchange occurs predominantly at topological defect sites, which often delineate distinct lattice configurations. Our results suggest that tau stabilizes longitudinal but destabilizes lateral tubulin dimer**–**dimer contacts. Through this change in anisotropy, tau effectively promotes the repair of structural defects by increasing their longitudinal mobility leading to their mutual annealing or removal from the lattice. Thus, tau does not only act as passive microtubule stabilizer but also facilitates the active elimination of lattice defects.

## Results

### Tau enhances tubulin incorporation into the microtubule lattice

We first investigated the influence of tau on tubulin incorporation into the microtubule lattice. We grew dynamic microtubules from GMPCPP-stabilized, surface-attached microtubule seeds using green-labeled GTP-tubulin (Fig. 1a, step I). To inhibit further microtubule tip dynamics, we capped microtubules with slowly hydrolyzable GMPCPP-tubulin (step II). We then incubated the microtubules with red-labeled GTP-tubulin in the presence of 0 nM, 0.5 nM or 20 nM human 2N4R tau for 15 min (step III). Tau covered the non-stabilized microtubule lattice sparsely and homogeneously, exhibiting diffusive motion and dynamic binding and unbinding on the timescale of a few seconds, as previously described^25–27^ (Fig. 1b, Fig. S1). We chose a relatively low free tubulin concentration of 8 µM to limit microtubule tip growth during the incorporation step. To visualize incorporated tubulin, we imaged the microtubules after washing out the free tubulin to reduce background fluorescence (step IV).

**Fig. 1:**
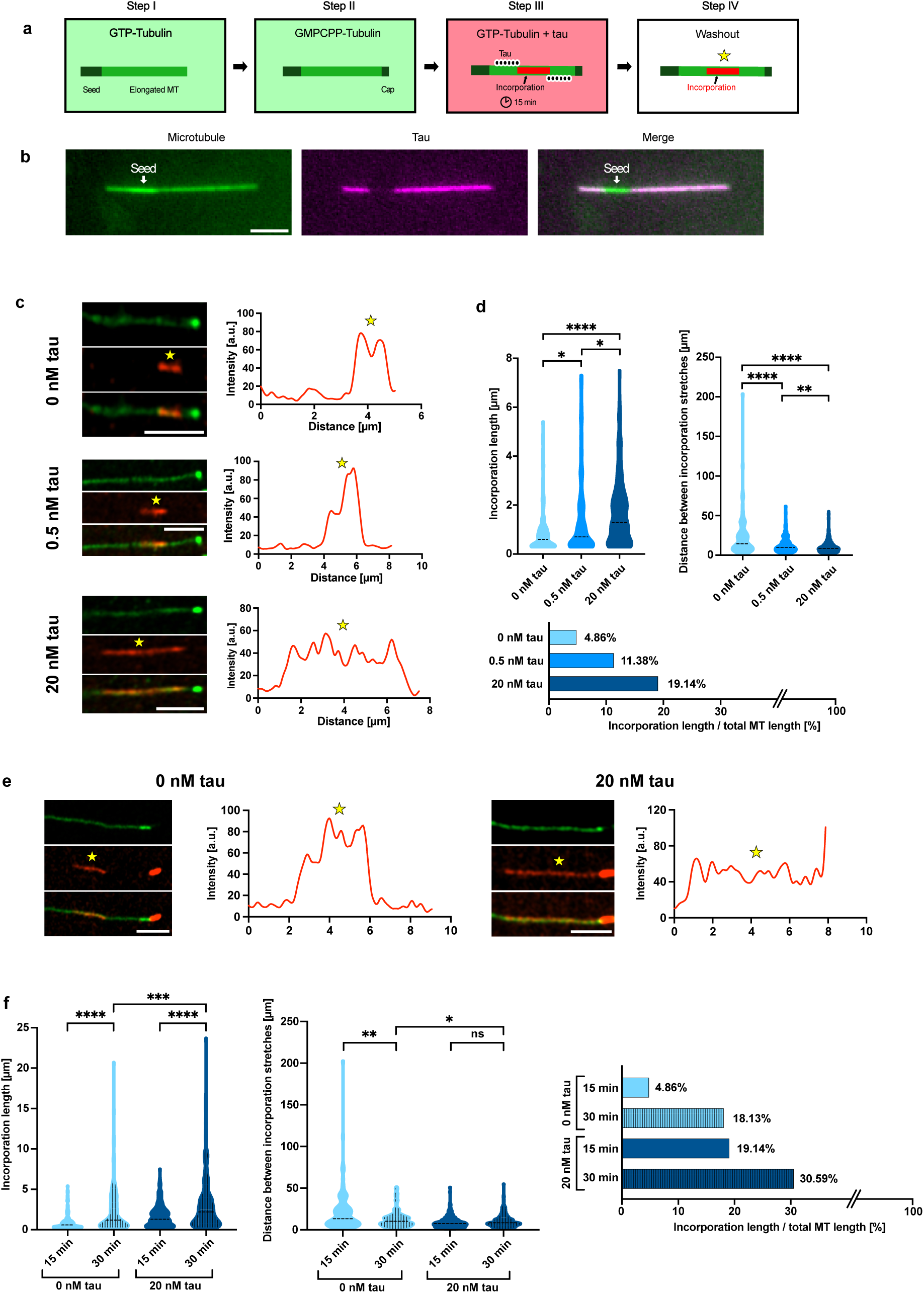
Tau increases tubulin incorporation into the microtubule lattice. **a.** Experimental setup to assess tubulin incorporation into the microtubule lattice in the presence of tau. First, dynamic microtubules were grown from GMPCPP-stabilized, surface-attached seeds in the presence of green-labeled GTP-tubulin (step I). Then, microtubules were capped with GMPCPP-tubulin to prevent further microtubule tip dynamics (step II). The microtubules were then incubated with 0 nM, 0.5 nM, or 20 nM unlabeled tau (black-white circles) and red-labeled GTP-tubulin for 15 min or 30 min (step III). The solution was replaced with buffer supplemented with taxol (step IV) to remove excessive background fluorescence, reveal tubulin incorporation sites (yellow star), and keep microtubules stable for imaging. **b.** 20 nM fluorescently labeled tau (magenta) homogeneously coat the microtubule lattice (green) before capping, with the exception of the GMPCPP-stabilized seed. Scale bar: 3 µm. **c.** Example images of microtubules (green) showing incorporation stretches after 15 min (red, stars) in the presence of 0 nM, 0.5 nM and 20 nM tau. The graphs show profile plots of the red channel along microtubules. Scale bars: 5 µm. **d.** Top left: The lengths of tubulin incorporation stretches increase in the presence of tau (> 2,200 µm microtubule length analyzed per condition; N=3 experiments per condition). Top right: The distances between incorporation stretches decrease in the presence of tau. Broken lines represent the median. Bottom: Tubulin incorporation is observed along 4.86% of the microtubule lattice at 0 nM tau, 11.38% at 0.5 nM tau and 19.14% at 20 nM tau. **e.** Example images with profile plots showing tubulin incorporation after 30 min in red (stars) and the lattice in green. Scale bars: 5 µm. **f.** Incorporation length (left) and distance between incorporation stretches (center) after 15 min vs. 30 min of incubation with red tubulin, for 0 nM and 20 nM tau. Broken lines represent the median. > 2,200 µm of microtubule length was analyzed in three independent experiments per condition. Tubulin incorporation was observed along 18.13% of the microtubule lattice for 0 nM tau and 30.59% for 20 nM tau (right).

When 20 nM tau was present during the tubulin-incorporation step, we noted extended tubulin incorporation stretches along microtubules, with median lengths of 1.22 µm compared to 0.67 µm in the absence of tau (Figs. 1c and 1d, top; Fig. S2; Methods). Furthermore, the median distance between individual incorporations decreased from 12.4 µm when no tau was present to 6.6 µm in the presence of 20 nM tau, implying an increased spatial frequency of incorporation events. These observations reveal a fourfold increase in tubulin incorporation in the presence of 20 nM tau compared to the control without tau (Fig. 1d, bottom).

Previously, we found that in the absence of other proteins, lattice dynamics constitute a continuous process^2^. To assess whether tau-stimulated tubulin incorporation also exhibits time-dependence, we extended the incubation time with red-labeled GTP-tubulin, supplemented with either 0 nM or 20 nM tau, to 30 min. Our observations revealed markedly longer tubulin incorporation stretches, occasionally spanning the majority of the microtubule length (Figs. 1e and 1f, left). While we also noted a higher spatial frequency of tubulin incorporation in the absence of tau after 30 min, the presence of 20 nM tau did not lead to a further increase in the spatial frequency of incorporation stretches (Fig. 1f, center). This suggests either a limitation in the number of available tubulin incorporation sites, reaching saturation more rapidly in the presence of tau, or a potential underestimation of these events due to extensive, overlapping incorporation stretches. Nevertheless, the trend of tau inducing a significant overall increase in tubulin incorporation remains consistent, with approx. 30% of the microtubule lattice showing tubulin incorporation in the presence of tau after 30 min (Fig. 1f, right). Taken together, tau significantly enhances tubulin incorporation into the microtubule lattice during an interaction time of up to 30 min.

### Tau reduces tubulin loss from the lattice

The observed marked increase in tubulin incorporation in the presence of tau may be ascribed to an elevated effective tubulin on-rate at the lattice, as demonstrated for the MAP CLASP^10^. Alternatively, it could stem from tau-induced tubulin loss from the lattice followed by subsequent incorporation of new tubulin (i.e., self-repair), a phenomenon observed e.g. for molecular motors^11–13^ and severing enzymes^14^.

To investigate the effect of tau on tubulin loss, we utilized tubulin incorporation as a read-out for preceding tubulin loss. We incubated capped green microtubules in tubulin-free buffer for a short duration (5 min), in the presence or absence of tau (Fig. 2a). Subsequently, we introduced red tubulin to allow for self-repair of the lattice sites which experienced tubulin loss. We observed decreased tubulin incorporation into microtubules coated with tau in the tubulin-free buffer (8.04% vs. 17.70% without tau), indicating a reduction in tubulin loss (Fig. 2b, c). Furthermore, we examined microtubule fracture as a result of potential tubulin loss. We grew capped microtubules as before (Fig. 2d, steps I-II) and then imaged them in the presence of 0 nM or 20 nM tau, but without free tubulin (steps III-IV). Fig. 2e shows an exemplary image sequence of a microtubule fracture event. Tau significantly slowed down microtubule fracture compared to the control without tau (Fig. 2f). Prior to fracture, the microtubule stretches with visible tubulin loss that indicate lattice damage became longer in the presence of 20 nM tau (Fig. 2g, Fig. S3). We note, however, that due to the longer time until fracture these data were recorded at later time points in the presence of tau than without tau. Jointly, our results indicate that tau reduces tubulin loss and thus, overall, stabilizes the microtubule lattice.

**Fig. 2:**
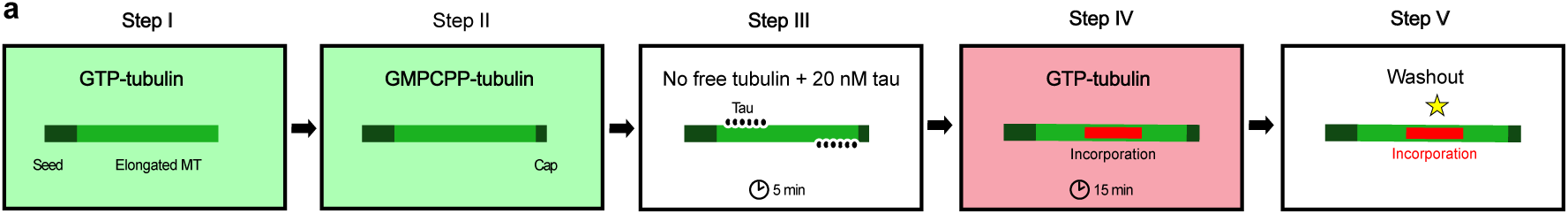
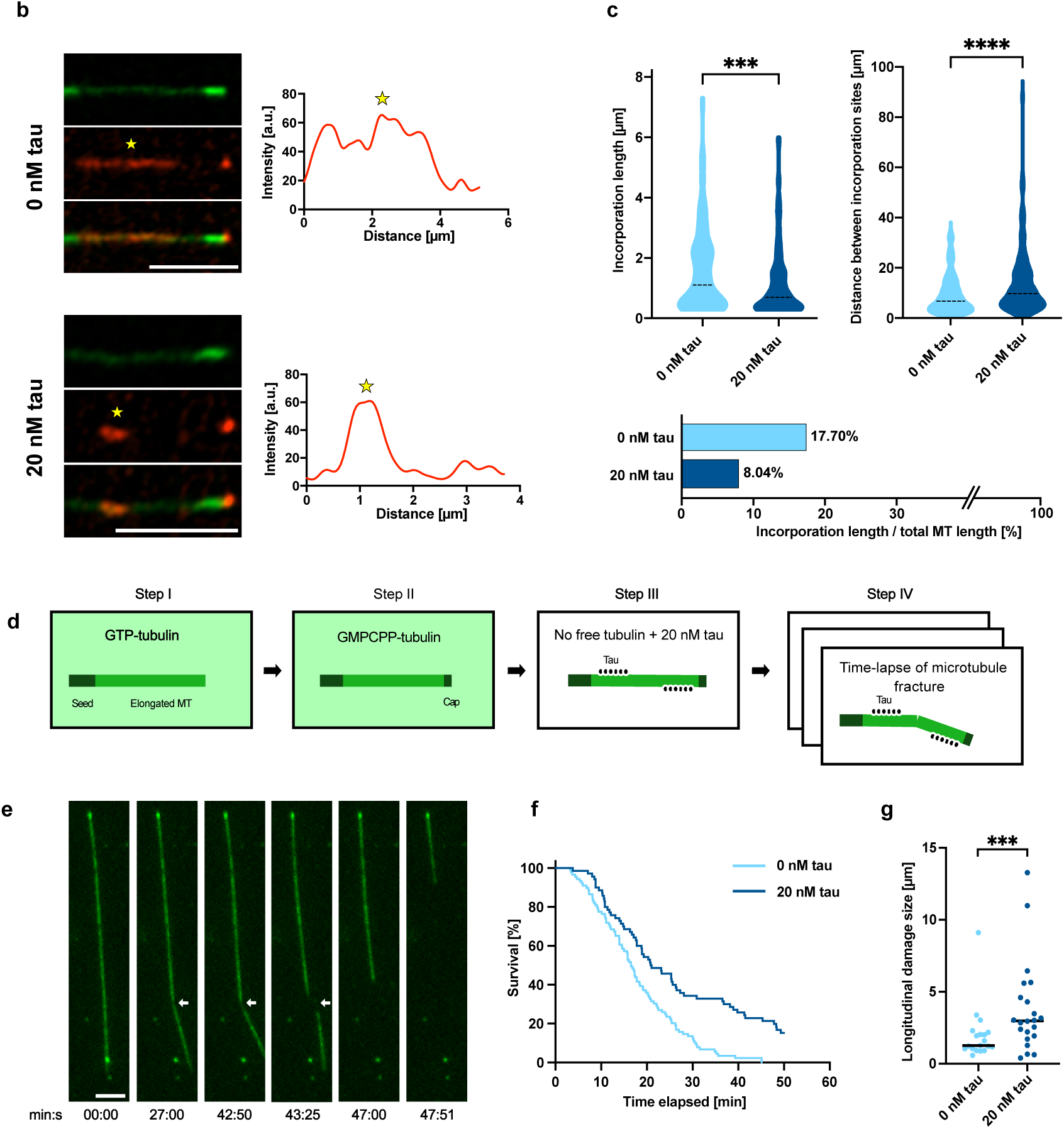
Tau reduces tubulin loss from the lattice. **a.** Experimental setup to confirm that tau reduces tubulin loss in the absence of free tubulin. Microtubules were grown with green labeled tubulin (step I) and capped (step II). They were then incubated with or without 20 nM tau in the absence of free tubulin for 5 min (step III), followed by a 15 min incubation step with red-labeled tubulin in the absence of tau (step IV). Microtubules were imaged after washout of free tubulin as before (step V). **b.** Example images showing tubulin incorporation (red, stars) into microtubules (green) after a 5 min incubation step without free tubulin. The profile plots show the fluorescence intensity along the microtubules in the red channel. Scale bars: 5 µm. **c.** Tubulin incorporation sites are shorter (top left) and more spaced (top right) in the presence of 20 nM tau compared to the control, leading to overall less tubulin incorporation (bottom). More than 1,700 µm of microtubule length was analyzed in three independent experiments per condition. **d.** Experimental setup to quantify microtubule fracture in the absence of free tubulin. Microtubules were grown with GTP-tubulin (step I) and capped with GMPCPP-tubulin (step II). Then, they were incubated with or without 20 nM tau in the absence of free tubulin (step III) and immediately imaged for 50 min (step IV). **e.** Image sequence showing a microtubule developing a damaged region, which is visible due to reduced green fluorescence (arrow) resulting from loss of tubulin from the lattice. The microtubule eventually broke along the softened region and disassembled. Scale bar: 3 µm. **f.** Survival curves showing that microtubule survival increases in the presence of tau. For 0 nM tau, 83 microtubules were analyzed in five independent experiments. For 20 nM tau, 65 microtubules were analyzed in four independent experiments. The differences in survival are statistically significant (p < 0.0001), based on a log-rank (Mantel-Cox) test. **g.** Lengths of the damaged lattice regions prior to fracture showing reduced fluorescence increased in the presence of tau. 19 and 22 fracture events were analyzed for 0 nM and 20 nM tau from five and four independent experiments, respectively.

### Tau stimulates tubulin exchange at defect sites

The lattice of a microtubule is never fully homogeneous; it inherently contains a variety of conformational changes^9^, resulting in the emergence of topological defects which can lead for example to protofilament transitions and seam dislocations along the length of a microtubule^2,6–8^. Such defects, often caused by monomeric tubulin vacancies associated with seam dislocations, weaken the microtubule lattice by disrupting local tubulin interactions. Tubulin incorporation at protofilament transitions and seam dislocations typically displaces rather than repairs these defects, as resolving them requires a conformational change along the microtubule. In fact, repair may only occur when a defect reaches the microtubule tip or merges with another defect of the same type. This contrasts with dimeric vacancies in an otherwise perfect lattice, where dimer incorporation can effectively heal the microtubule.

We previously demonstrated that defects are likely sites of lattice dynamics^2,3^, prompting us to investigate whether tau-enhanced tubulin incorporation also occurs preferentially at lattice defect sites. Towards this end, we conducted tubulin incorporation experiments in the presence of tau and subsequently imaged microtubules in tubulin-free buffer until fracture (Fig. 3a). Indeed, our observations reveal that while incorporation stretches covered only about 20% of the lattice, around 40% of fracture events occurred at these stretches (Fig. 3b). This suggests that tau-enhanced tubulin incorporation also occurs preferentially at preexisting lattice defect sites, which are not repaired but only translocated. Thus, the incorporation stretches likely mark the traces of the translocating defects. In line with this picture, we frequently observed the disappearance of the incorporation stretch before microtubule fracture (Fig. 3c).

**Fig. 3:**
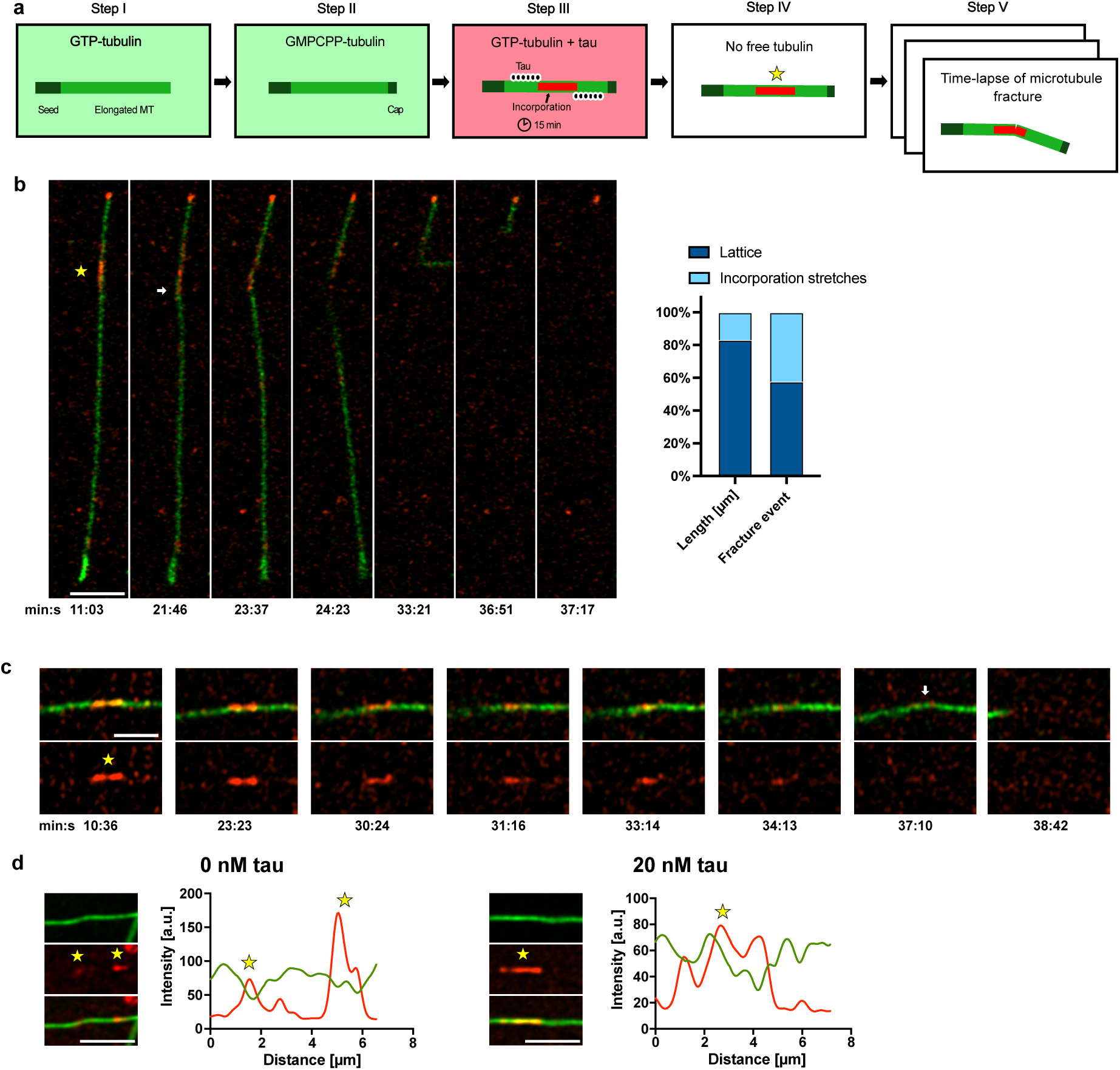
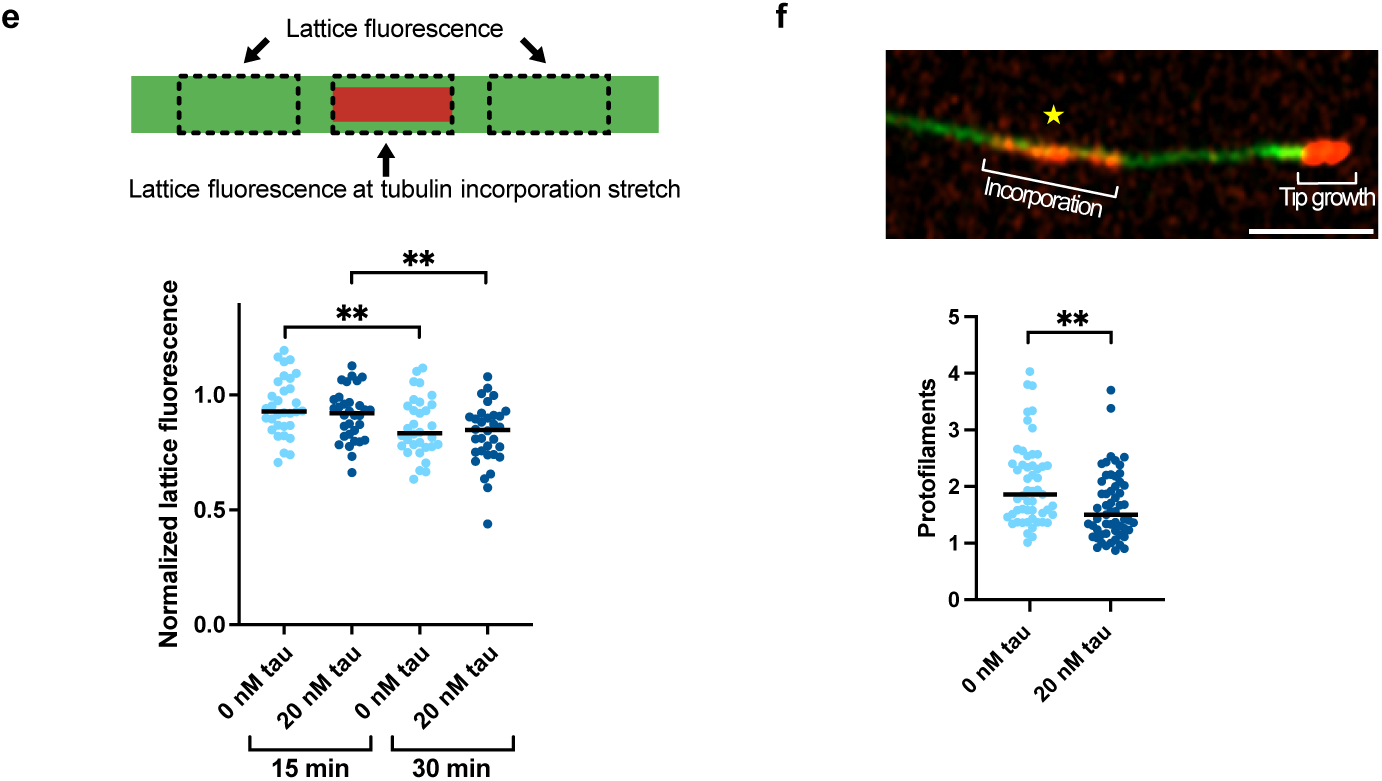
Tau stimulates tubulin exchange. **a.** Experimental setup to visualize microtubule fracture after tubulin incorporation. Microtubules were grown with green-labeled tubulin (step I) and capped (step II) before incubation with 20 nM tau and red-labeled tubulin (step III). Then, the solution was replaced with buffer without taxol (step IV) and immediately imaged for 50 min (step V). **b.** Left: Example images showing a microtubule that developed a kink (arrow) at an incorporation stretch (red, star) before fracture at this site. Scale bar: 3 µm. Right: Microtubule fracture occurred more frequently at tubulin incorporation sites than is to be expected based on incorporation lengths compared to the total microtubule length. For 20 nM tau, 26 microtubules were analyzed in three independent experiments. **c.** Example images showing that the incorporation stretch (red, star) disintegrated before the microtubule developed a kink (arrow) and disassembled. Scale bar: 3 µm. **d.** Example images of tubulin incorporation after 30 min (red, stars) into the microtubule lattice (green), averaged over at least five consecutive time points, with fluorescence profile plots in the absence and presence of 20 nM tau. Scale bars: 3 µm. **e.** Lattice fluorescence at incorporation stretches, normalized to the fluorescence of the lattice next to the incorporation stretches. Black lines represent the median. 32 incorporation sites from three different experiments were analyzed per condition. **f.** Estimation of the number of protofilaments constituting an incorporation stretch. Top: Example image of an incorporation stretch (star) and the microtubule tip consisting of red-labeled tubulin, which was used as a reference to estimate the number of protofilaments present at the incorporation stretch. Scale bar: 3 µm. Bottom: Number of protofilaments constituting an incorporation stretch in the absence and presence of tau. 53 and 57 incorporation sites were analyzed from three different experiments for 0 nM and 20 nM tau, respectively. Black lines represent the median.

Does free tubulin from solution replace dimers in the microtubule or is it ‘added’ to the pre-existing lattice? When plotting the fluorescence intensity along microtubules, the original (green) lattice signal often appeared to slightly decrease at stretches of newly incorporated (red) tubulin, both in the presence and absence of tau (Fig. 3d). Because the signal is relatively noisy, we integrated the original (green) lattice signal over the length of the incorporation stretch and compared it to the original (green) lattice signal next to this stretch (normalized lattice fluorescence, Fig. 3e, top). We found that, on average, the original (green) lattice intensity was slightly lower at the incorporation sites (represented by averaged normalized lattice fluorescence values <1), with a more pronounced reduction after 30 min than after 15 min (Fig. 3e, bottom). These observations show that the incorporated tubulin does indeed replace tubulin from the original lattice. Due to the noise in our measurements of the normalized lattice fluorescence, we cannot detect a possible influence of tau on the number of protofilaments over which tubulin is replaced. We therefore aimed to estimate the number of protofilaments on which tubulin is incorporated by normalizing the fluorescence intensity of the red-labeled incorporation stretches by the fluorescence intensity of a red-labeled microtubule tip growing with about 13 protofilaments in the presence of GTP (Fig. 3f, 30 min incubation time with red tubulin). We found that the incorporation stretches often consisted of one to two protofilaments, occasionally reaching a maximum of up to four, indicating that only a small portion of the lattice is exchanged. Interestingly, the number of exchanged protofilaments was lower in the presence of tau.

### Tau increases lattice anisotropy

At first glance, our results seem unexpected, as the fracture experiments in the absence of free tubulin point to a stabilizing effect of tau, whereas the tau-enhanced exchange of tubulin in the vicinity of lattice defects necessarily requires an accelerated tubulin dissociation. This apparent contradiction raises the question how tau acts on the lattice, specifically regarding the bonds between neighboring tubulin dimers.

A recent high-resolution cryo-electron microscopy study revealed that tau’s microtubule-binding domains (MTBDs) bind along individual protofilaments, each bridging an intra- and an inter-dimer contact potentially affecting longitudinal tubulin interactions^24^. To test this, we performed coarse-grained molecular dynamics (MD) simulations of a tau MTBD on three longitudinally connected tubulin monomers and used umbrella sampling to compute the free energy profile along the dissociation of a longitudinal dimer–dimer contact (Fig. 4a). Here, the free energy minimum at a distance of approximately 4 nm corresponds roughly to the known equilibrium state^28–31^. As shown by the difference between the dark and light blue curves in Fig. 4a, the binding free energy is lower in the presence of tau than in its absence. Although the approximations underlying our coarse-grained simulations do not permit a precise quantification of the difference in binding free energies with and without tau, they strongly suggest that tau stabilizes longitudinal dimer–dimer contacts, likely facilitating overall microtubule stability.

**Fig. 4:**
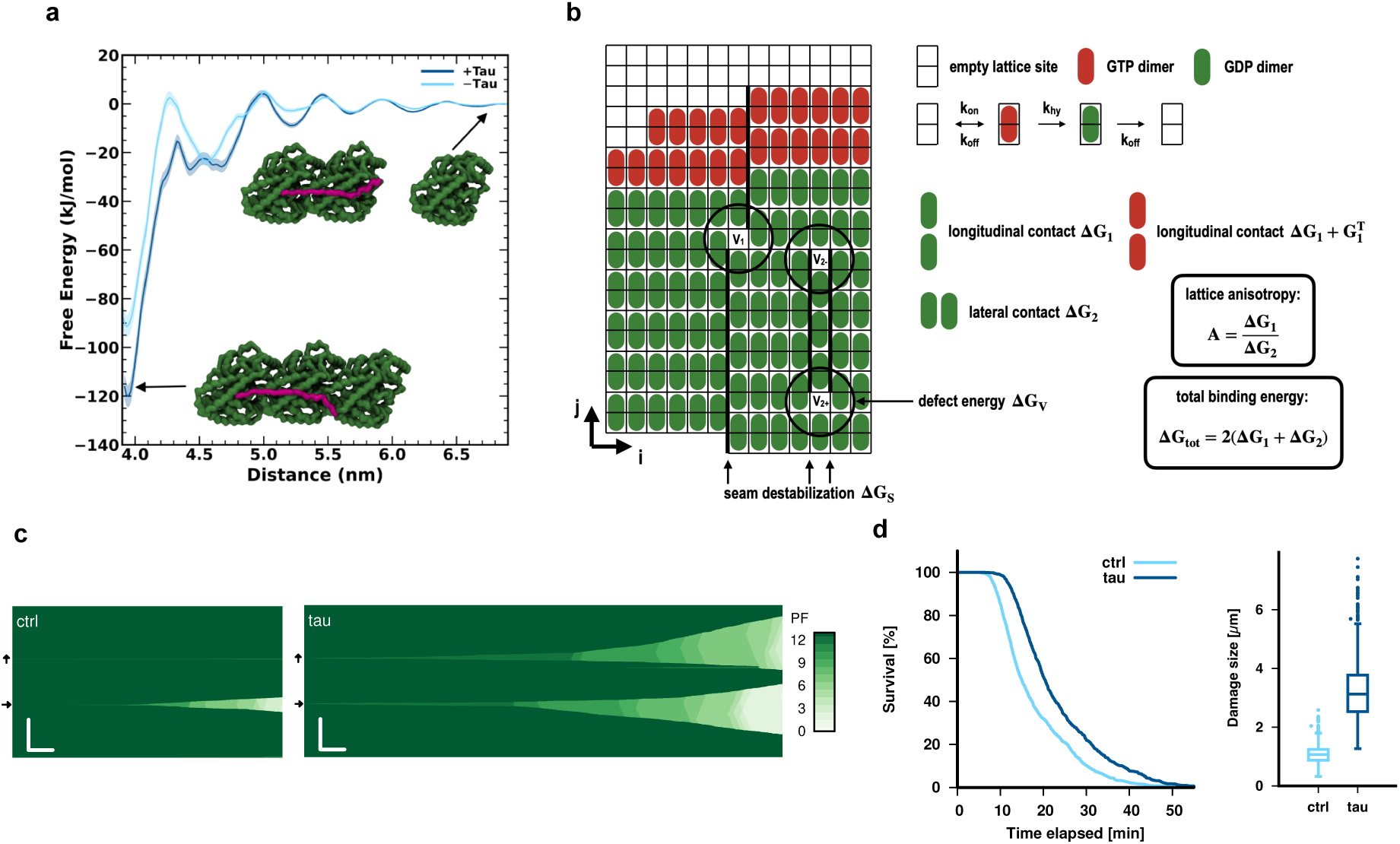
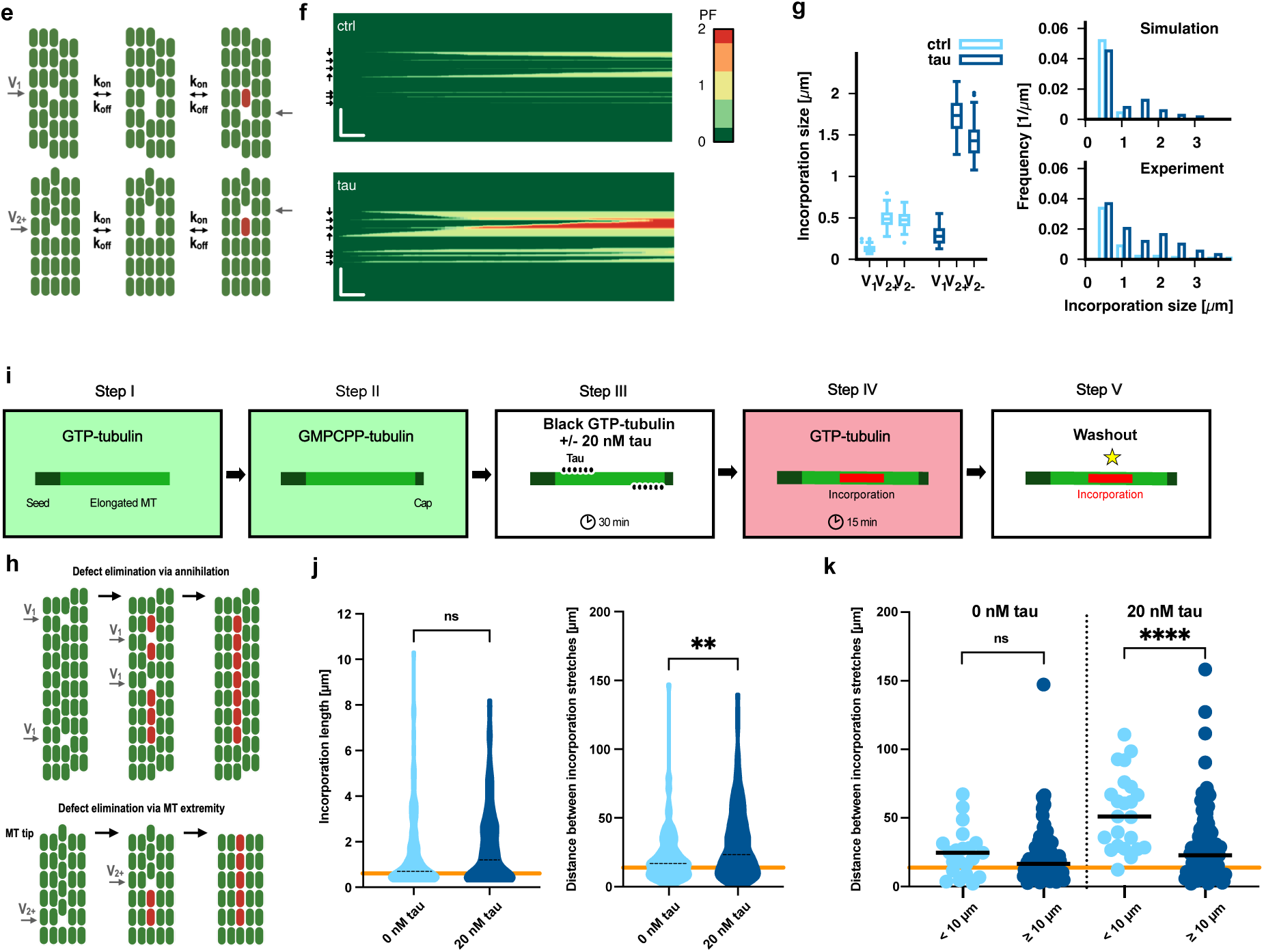
Model of the effect of tau on microtubule lattice dynamics. **a.** Tau stabilizes longitudinal dimer–dimer contacts. Free energy profiles for tubulin dissociation in the presence (dark blue curve) and absence (light blue curve) of tau (magenta) as a function of the center-of-mass distance between adjacent monomers. In the presence of tau, the binding affinity along longitudinal dissociation of tubulin dimer and monomer is stronger than in the absence of tau. Shaded areas represent the standard error of the mean. The inset shows the structure of three tubulin monomers (green, top view) with a single monomer depicted in both the associated and dissociated states from the microtubule-tau complex. **b.** Schematic representation of the kinetic Monte Carlo model for the microtubule lattice. The microtubule is modeled as the canonical 13 protofilament three-start helix structure on the scale of the monomer, such that the dynamics of monomer vacancies (labeled V_1_, V_2+_, and V_2-_) can be investigated. V_2+_ and V_2-_ refer to the orientation of the newly created seams towards the microtubule plus-end and minus-end, respectively. Shown are also the three possible kinetic transitions with their respective rate constants (k_on_: GTP-dimer attachment, k_off_ GTP or GDP dimer detachment, k_hy_ GTP dimer hydrolysis) and the relevant energetic contributions which govern the kinetics of dimer detachment within Kramer’s theory. **c.** Simulation of microtubule fracture of end-stabilized microtubules. Kymographs of a representative lattice configuration with one V_1_- and one V_2+_-type defect marked by → and ↑ arrows, respectively, along a 10 µm long microtubule. Scale bars: 2 µm vertical, 1 min horizontal. The color code indicates the number of intact protofilaments. Parameters are ΔG^S^=0.5 kT, and ΔG_tot_=-36 kT, A=1.5 (control) and ΔG_tot_=-36.2 kT, A=2.1 (tau) and as indicated in Table S2. **d.** Left: Simulated survival curves of end-stabilized microtubules in the absence of free tubulin, in the presence (dark blue) and absence (light blue) of tau. The curves show the survival of 1000 microtubules with randomly placed defects (frequency 0.15 𝜇m^-1^). Right: Simulated longitudinal damage size, corresponding to the loss of all 13 protofilaments. Dark blue denotes the simulations with tau, light blue the control case without tau. **e.** Schematic representation of defect motion via dimer detachment and attachment. **f.** Simulation of dimer incorporation dynamics into end-stabilized microtubules. Kymographs of a representative lattice configuration with V_1_- as well as V_2+_- and V_2-_-type defects marked by →, ↑ and ↓ arrows, respectively, along a 10 µm long microtubule. Scale bars: 2 µm vertical, 1 min horizontal. The color code indicates the number of exchanged protofilaments. Parameters are ΔG_S_=0.5 kT, and ΔG_tot_=-36 kT, A=1.5 (control) and ΔG_tot_=-36.2 kT, A=2.1 (tau) with ΔG_V_=1.25 kT and as indicated in Table S2. To compare the simulated results with experiments, the longitudinal positions of incorporated dimers were convoluted with a point spread function as described in the SI. **g.** Length and frequency of incorporation stretches. Left: Incorporation length depending on the defect type (V_1_, V_2+_, V_2-_) in the presence of tau (dark blue) compared to the control (light blue) after 15 min. Right top: Simulated distribution of (visible) incorporation sizes after 15 min in the presence (dark blue) and absence (light blue) of tau for 10 µm long microtubule with randomly placed defects (frequency 0.15 𝜇m^-1^) showing a bimodal distribution in the presence of tau. The frequency of visible incorporations are 0.056 𝜇m^-1^ (control) and 0.076 𝜇m^-1^ (with tau). To compare the simulated results with experiments, the longitudinal positions of incorporated dimers were convoluted with a point spread function as described in the SI. Right bottom: Experimental distribution of visible incorporation sizes. The data correspond to the data shown in Fig. 1d. **h.** Exemplary representation of the elimination of two V_1_ lattice defects via annealing (top) and a V_2+_ defect via motion up to the microtubule extremity (bottom). **i.** Experimental setup to visualize tubulin incorporation after an initial 30 min tubulin incorporation step in the presence or absence of tau. Microtubules were grown with green-labeled tubulin (step I) and capped (step II) before incubation with unlabeled tubulin in the presence or absence of 20 nM tau for 30 min (step III). This initial incorporation step was followed by an additional 15 min incorporation step in the presence of red-labeled tubulin and in the absence of tau (step IV) before washout and imaging (step V). **j.** Incorporation lengths (left) and distances between incorporation stretches (right). Broken lines represent the median. Orange lines represent the median values from Fig. 1d (incorporation length and distance between incorporations for 0 nM tau) as reference, respectively. The resulting tubulin incorporation lengths are comparable between the samples, but the control sample shows slightly more frequent tubulin incorporations (> 3,000 µm of microtubule length was analyzed from three independent experiments per condition). **k.** Distances between tubulin incorporation stretches from the graph on the left were binned according to microtubule length, showing a marked increase for short (0-10 µm) microtubules in the presence of tau. Black lines represent the median. The orange line represents the median from Fig. 1d (distance between incorporation stretches for 0 nM tau) as reference.

The observed tau-enhanced exchange of tubulin requires, however, lattice destabilization, which needs to originate from weakened lateral dimer–dimer contacts. Weakened lateral interactions would increase the lattice anisotropy (ratio of longitudinal to lateral tubulin bond energies), in line with our experimental observation of a predominantly longitudinal spreading of tubulin loss prior to fracture when tau is present (Fig. 2g): the higher the lattice anisotropy, the more likely it is to lose a dimer that misses a longitudinal bond compared to a lateral bond, favoring pronounced longitudinal spreading of damage sites along the lattice. Moreover, such increase in anisotropy would explain why we observe less lateral spreading of exchanged tubulin (i.e., incorporation over fewer protofilaments) in the presence of tau (Fig. 3f).

### Numerical modeling captures the dual effect of tau on lattice dynamics

As proof of principle, we developed a semi-quantitative kinetic Monte Carlo model for the microtubule lattice^2,28,32,33^ that captures relevant time and length scales with a minimum set of parameters. Most importantly, we model the dynamics of monomer vacancies that lead to seam dislocations, which are the most frequent defects in vitro and which may lead to the creation of additional A-lattice contacts (multi-seam structures^8^). Based on the available structural information^8^, we introduced two distinct types of monomer vacancies, denoted as V_1_ and V_2_ (Fig. 4b). For V_1_-type defects, an existing A-lattice seam undergoes a lateral shift by one protofilament. In contrast, V_2_-type defects generate two additional A-lattice seams within a B-lattice, extending either towards the plus- or minus-end of the microtubule.

To model the lattice dynamics at seam dislocations, we made the following assumptions (see Fig. 4b; Fig. S4, refer to the model description in the SI for further details): 1) Since A-lattice contacts lead to an increased lateral opening between protofilaments^34^ and microtubules exhibiting ectopic A-lattice seams depolymerize at an accelerated rate_35_, we assign an additional weak destabilization ΔG_S_>0 to lateral A-lattice contacts compared to B-lattice contacts. 2) Seam dislocations induce a sharp shift in the lattice structure on the scale of a few tubulin dimers. In solid mechanics, it is well established that defects distort the lattice structure of highly ordered materials and are associated with a nonlocal stress field for various systems ranging from metals to colloidal crystals^36,37^. Therefore, we assign a small destabilization ΔG_V_>0 to the dimers around the seam dislocations that we limit to the nearest neighbors for simplicity and which we refer to as ‘defect energy’. 3) The effect of tau is included by effectively modulating the binding energy (ΔG_tot_) and lattice anisotropy compared to the microtubule lattice without tau. Note that we assume that tau only affects the lateral and longitudinal dimer-dimer contacts (point 3 above) while leaving the lateral A-lattice destabilization and the defect energy unaffected (points 1 and 2 above). To identify the experimentally relevant lattice parameters, we reproduced the fracture behavior observed in our experiments (Fig. 4c, d; Fig. S5): we found that an increase in the lattice anisotropy from 1.5 in the absence of tau to 2.1 in the presence of tau, along with a slight net stabilization of the microtubule lattice (−0.2 kT), reproduced the increase in fracture time (Fig. 2f) and the longitudinal damage spreading in the presence of tau (Fig. 2g).

Next, we used our model to investigate tubulin incorporation at defect sites. V_1_ defects are symmetric (Fig. 4e, top). As continued tubulin loss and incorporation occurs, they move mainly diffusively along a protofilament (Fig. 4f and Fig. S10). In contrast, V_2_ defects exhibit a mainly ballistic motion due to their longitudinal asymmetry (Fig. 4e, bottom; Fig. 4f and Fig. S10). Thus, V_2_ defects generally produce longer tubulin incorporation stretches (kymographs in Fig. 4f and Fig. 4g, left). Note that at accelerated defect dynamics in the presence of tau, V_1_ defects move preferentially towards the microtubule plus-end, while the minus-end directed motion of V_2-_ defects slows down compared to V_2+_ defects (Fig. 4g left; Fig. S10). Both effects result from the different mechanisms of GTP-hydrolysis of the tubulin 𝛽-subunit at the microtubule plus- and minus-end. The first peak in the incorporation length visible in the presence of tau at around 0.5 µm can be attributed to V_1_ defects, while the second peak at around 1.5 µm stems mostly from faster moving V_2_ defects (Fig. 4g, right top). This bimodal distribution is also visible in the experimental data (Fig. 4g, right bottom). Taken together, tau accelerates lattice dynamics for both V_1_ and V_2_ defects, leading to longer tubulin incorporation stretches than in the control without tau.

Our simple model recapitulates the dual effect of tau: overall, it leads to a stabilization of the microtubule lattice, while it also increases lattice dynamics at both V_1_ and V_2_ defect sites. This allows the interesting prediction that tau accelerates defect motion along a protofilament. As a consequence, such enhanced mobility would increase the probability for defects to meet and anneal in a given time interval (Fig. 4h, top). Moreover, ballistically moving V_2_ defects may also propagate until they reach the end of the non-stabilized microtubule, leaving behind a repaired trace (Fig. 4h, bottom). To test this prediction experimentally, we incubated green capped microtubules with unlabeled tubulin in the presence of 0 nM or 20 nM tau for 30 min to allow defects to anneal or move to the end of the non-stabilized lattice (Fig. 4i, steps I-III; refer to Fig. 1f). We then incubated the microtubules with red-labeled tubulin in the absence of tau for 15 min (step IV) before washout (step V). While we found no significant difference in the lengths of incorporation sites, the tau-incubated microtubules showed a significant decrease in the spatial frequency of incorporations (Fig. 4j). Since the chance for defects to travel to the end of the non-stabilized lattice is higher the shorter a microtubule is, we separated our results according to the lengths of the non-stabilized microtubule parts. Indeed, we found a pronounced effect for lengths up to 10 µm in the presence of tau, where the median distance between incorporation sites increased more than two-fold from 20.3 µm in the control (without tau) to 51.1 µm (Fig. 4k).

## Discussion

In summary, we experimentally found that tau has a dual effect on microtubule lattice dynamics: it accelerates tubulin turnover distant from the tips, while overall stabilizing the lattice. We propose a numerical model that recapitulates our experimental findings. We opted for a simple model that explains the essential trends we observed in our experiments without aiming for a precise quantitative match, thus avoiding the introduction of additional parameters that cannot be defined with current experimental data.

In our model, tau increases the lattice anisotropy by strengthening longitudinal bonds, as supported by our molecular dynamics simulations, and weakening lateral bonds (Fig. 5). Tau also slightly increases the overall bond strength of tubulin dimers in the lattice. These (weak) modifications in tubulin bond energies are sufficient to explain the dual effect of tau and align with the previous descriptions of tau slowing down microtubule depolymerization^20^ and may also account for the previously observed increase in microtubule stiffness in the presence of tau^38–42^.

**Fig. 5:**
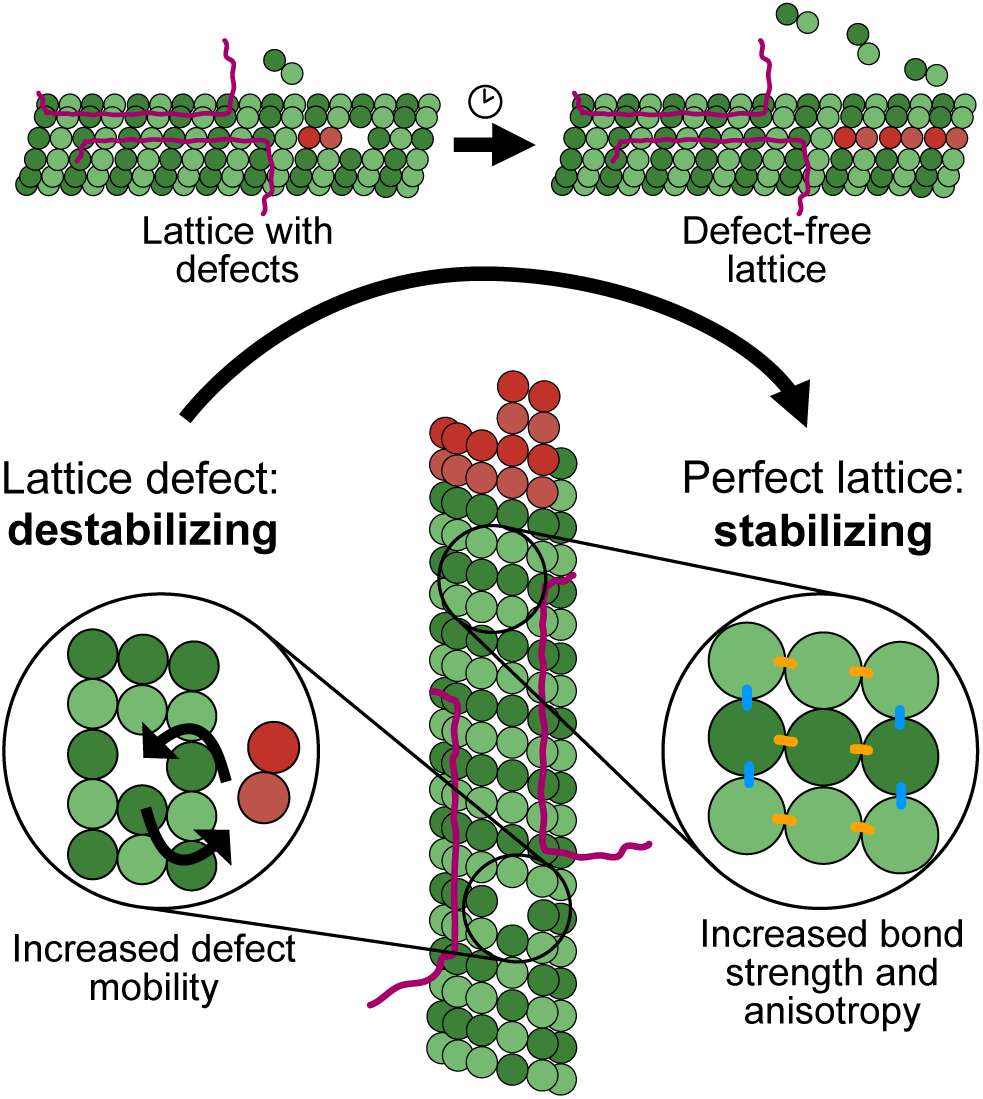
Conceptual model of the dual effect of tau on microtubule lattice dynamics. Based on our experimental results and numerical simulations, we propose a model in which tau (magenta) increases the anisotropy of tubulin bond energies in the lattice by strengthening longitudinal bonds (blue) and weakening lateral bonds (orange). Overall, tau has a stabilizing effect on the intact lattice that is consistent with the previously observed reduction in microtubule depolymerization speed. At lattice defects, tau increases the exchange of tubulin dimers; the above-mentioned changes in tubulin bond strengths are sufficient to explain this effect. In this way, tau accelerates the mobility of defects, driving them to the microtubule end or to mutual annihilation and thus contributing to healing lattice defects.

Interestingly, a recent study reported that tau increases the number of lattice defects when present during microtubule polymerization^23^. Although the origin of this effect is not yet clear, it suggests that tau plays different roles during microtubule polymerization and after microtubule growth. During polymerization, tau appears to stabilize alternative lattice configurations, whereas, after microtubule growth, our observations indicate that tau accelerates the dynamics of lattice defects and ultimately facilitates their removal. Once the lattice defects are removed, tau functions as a genuine microtubule stabilizer. Thus, tau may be considered a versatile caretaker of the microtubule lattice, combining both passively stabilizing and actively modulating functionalities.

## Methods

### Molecular cloning, expression and purification of Tau

The gene coding for Tau2N4R was amplified by polymerase chain reaction using the primers flanked with the restriction sites NotI and AscI respectively at 5’ (AATAATAACATGCGGCCGCAATGGCTGAGCCCCGCC) and 3’ (AATAATAACATGGCGCGCCCAAACCCTGCTTGGCCAG) ends. The resulting product was inserted in the FlexiBac^43^ vector backbone with either c-terminal monoGFP followed by 3C protease cleavage site and 6xHistadine tag or without the fluorescent monoGFP tag. The construct sequences were then confirmed by digestion and sequencing. Plasmid vectors with tau2n4r gene was then co-transfected with a replication-defective bacmid DNA into Sf9 cells. Homologous recombination between flanking sequences in both piece of DNA introduces the gene of interest into the viral genome and rescues viral replication and subsequent amplification in Sf9 insect cells. For protein expression SF9 cells were infected with recombinant passage 2 viruses with a ratio of 1:100 (v/v) and harvested 72 hours post infection by centrifugation at 500 x g for 10 min. Harvested cells were resuspended in the lysis buffer (25 mM HEPES, 150 mM KCl, 5% glycerol, 0.1% Tween20 (v/v), 20 mM Imidazole, 1 mM TCEP and 1 x protease inhibitor cocktail (cocktail III Merck calbiochem, 535140). Cells were then lysed using an Emulsiflex (Emulsiflex-C5, Avestin). All further steps were then performed at 4°C or ice. The lysate was then spun at 186,000 x g for 45 min. The supernatant was filtered through a 0.45 μm syringe filter and passed through a 1 ml His-trap column (29051021, Cytiva) pre-equilibrated with 10 coulmn volume of equilibration buffer (50 mM HEPES, 300 mM KCl, 5% glycerol, 0.2% Tween20 (v/v), 20 mM Imidazole, 1 mM TCEP pH 7.2). The protein was then eluted by passing the elution buffer (equilibration buffer supplemented with 200 mM Imidazole). 6x His tag was then removed by incubating the eluted fractions with PreScission protease (3C HRV protease, 1:100, 1µg enzyme/100 µg of protein, overnight at 4°C). Protease treated protein fractions were then concentrated using 30 KDa MWCO Amicon spin filters with buffer exchanged to equilibration buffer. The protein of interest was further purified by size exclusion chromatography (SEC) using Superdex™ 200 Increase (28990944, Cytiva) column with and ÄKTA Pure chromatography system (GE Healthcare) in storage buffer (25 mM HEPES, 150 mM KCl, 5% glycerol, 0.1% Tween20 (v/v), 1 mM TCEP pH7.2). Collected peak fractions were concentrated to using Amicon Ultra 30K (Millipore), aliquoted and then snap-frozen in liquid nitrogen and stored at −80°C. SDS-PAGE was used to determine the purity of isolated proteins and for protein concentration estimation by running serial dilutions of BSA (1 mg/ml) along-side appropriate dilutions of the proteins on a 4– 12% BisTris SDS-PAGE precast gel in MOPS buffer (Life Technologies), stained with Coomassie for 60 min and de-stained in distilled water. Stained gels were imaged in an imaging station (c300, Azure Biosystems). The integrated intensity of the protein bands of interest was quantified using the “Gels” tool in Fiji^44^. Linear fitting of the integrated intensity vs. concentration of BSA provided a calibration curve which was then used to estimate the concentration of desired proteins.

### Tubulin purification and labeling

Tubulin was purified from fresh bovine brain by three cycles of temperature-dependent assembly and disassembly in Brinkley Buffer 80 (BRB80 buffer; BRB buffer: 80 mM PIPES, pH 6.8, 1 mM EGTA and 1 mM MgCl_2_ plus 1 mM GTP). MAP-free neurotubulin was purified by using low and high salt buffer (High Molarity PIPES Buffer) in 1M PIPES, pH 6.9, supplemented with KOH, 10 mM MgCl_2_, 20 mM EGTA and subsequent cation-exchange chromatography (EMD SO, 650 M, Merck) in 50 mM PIPES, pH 6.8, supplemented with 1 mM MgCl^2^ and 1 mM EGTA. Purified tubulin was obtained after, in total, three cycles of polymerization and depolymerization. Fluorescent tubulin (ATTO-488 and ATTO-565-labeled tubulin) and biotinylated tubulin were prepared as previously described^45^: Microtubules from neurotubulin were polymerized at 37 °C for 30 min and layered onto cushions of 0.1 M NaHEPES, pH 8.6, 1 mM MgCl_2_, 1 mM EGTA, 60% v/v glycerol, and sedimented by high centrifugation at 37 °C. Then microtubules were resuspended in 0.1 M NaHEPES, pH 8.6, 1 mM MgCl_2_, 1 mM EGTA, 40% v/v glycerol and labeled by adding 1/10 volume 100 mM NHS-ATTO (ATTO Tec), or NHS-LC-LC-biotin (EZ-link, Thermo) for 10 min at 37 °C. The labeling reaction was stopped using two volumes of 2x BRB80, containing 100 mM potassium glutamate and 40% v/v glycerol, and then microtubules were sedimented onto cushions of BRB80 supplemented with 60% glycerol. Microtubules were resuspended in BRB80, and an additional cycle of polymerization and depolymerization was performed before use.

### Cover glass treatment

The micropatterning technique was adapted from Portran et al.^46^. Cover glasses were cleaned by successive chemical treatments: 30 minutes in acetone, 15 minutes in ethanol (96%), rinsing in ultrapure water, 2 hours in Hellmanex III (2% in water, Hellmanex), and rinsing in ultrapure water. Cover glasses were air dried and incubated for three days in a solution of tri-ethoxysilane-PEG (30 kDa, Creative PEGWorks) or a 1:10 mix of tri-ethoxy-silane-PEG-biotin and tri-ethoxy-silane-PEG at 1 mg ml^−1^ in ethanol 96% and 0.02% HCl, with gentle agitation at room temperature. Cover glasses were then successively washed in ethanol and ultrapure water, air dried, and stored at 4°C.

### Microtubule growth, capping, and tubulin incorporation

Microtubule seeds were prepared at 10 μM tubulin concentration (30% ATTO-488-labeled or ATTO-565-labeled tubulin and 70% biotinylated tubulin) in BRB80 supplemented with 0.5 mM GMPcPP at 37 °C for 1 hour. The seeds were incubated with 50 μM Paclitaxel (Merck) at room temperature for 30 minutes and then sedimented at 156,000 x g at 25°C and resuspended in BRB80 supplemented with 0.5 mM GMPcPP and 50 μM Paclitaxel. Seeds were stored in liquid nitrogen and quickly warmed to 37 °C before use.

A flow cell chamber with an approximate volume of 40 μl with an entry and exit site was constructed with double-sided adhesive tape with a glass coverslip functionalized and passivated with SiPEG (top) or SiPEG and SiPEG-Biotin (9:1, bottom). The flow chamber was perfused for 30 seconds with neutravidin (50 μg ml^−1^ in 1x BRB80; Fisher Scientific), passivated for 30 seconds with PLL-g-PEG (Pll 20 K-G35-PEG2K, Jenkam Technology) at 0.1 mg ml^−1^ in 10 mM Na-HEPES (pH = 7.4), and washed again with 1x BRB80. Microtubule seeds were flushed into the chamber at high flow rate to ensure parallel orientation of the seeds to the flow direction. Non-attached seeds were washed out immediately using BRB80 supplemented with 1 mg/ml Casein.

Seeds were elongated with a mix containing 11 μM of tubulin (5% to 20% labeled, green fluorescent) in 0.7x BRB80 and 0.38x MAP Buffer (500mM Phosphate Buffer, 1M KCl, 10mM DTT, pH 7.9) supplemented with 1 mM GTP, an oxygen scavenger cocktail (22 mM DTT, 1.2 mg ml^−1^ glucose, 8 μg ml^−1^ catalase and 40 μg ml^−1^ glucose oxidase), 1 mg/ml casein and 0.033% (w/v) methyl cellulose (1,500 cP, Sigma) at 37°C. GMPcPP caps were grown by substituting in the before-mentioned buffer GTP with 0.5 mM GMPcPP (Jena Bioscience) and using 3 μM tubulin (100% labeled with ATTO-488) at 37°C. Capping extends the microtubule lifetime and allows for the examination of lattice turnover over a period of several minutes despite the dynamic instability *in vitro*. For incorporation experiments, the same buffer as for seed elongation was used, supplemented with 8 μM tubulin (100% labeled with ATTO-565) along with 0 nM, 0.5 nM, or 20 nM 2N4R-GFP tau or unlabeled 2N4R tau diluted in 1x BRB80. Microtubules were incubated in this buffer for 15 minutes or 30 minutes at 37 °C before replacing it with a washing buffer (with the same buffer as for seed elongation with 50µM paclitaxel and without free tubulin) for imaging. For fracture experiments, a buffer containing the same supplements but without taxol and free tubulin was used. For tubulin incorporation along with fracture experiments, first incorporation was performed as mentioned above both in the presence and absence of tau and subsequently imaged microtubules in tubulin-free buffer until fracture as discussed earlier.

### Imaging and image analysis

Microtubules were visualized using a point scanning confocal microscope (Zeiss LSM 900) with Airyscan 2 and Axiocam 705 camera. The microscope stage was kept at 37°C through a cage incubator (PECON). Time-lapses were recorded using ZenBlue software (version 3.2).

Microtubule fracture experiments were performed and visualized on an objective-based orbital TIRF microscope (Nikon Eclipse Ti2, modified by ViSitron Systems) and an EMCCD Camera (Andor iXon Life) at minimal laser intensity. The microscope stage was kept at 37 °C using a warm stage controller (OkoLabs). Time-lapse recording was performed using VisiView software (version 6.0).

To visualize incorporation, images were typically taken every 0.5 seconds, and 5 images were overlaid and averaged. Line scans of the red fluorescence intensity along the microtubule (Fig. 1c) were used to identify the stretches of incorporated tubulin. Stretches of tubulin incorporation were identified as corresponding to zones of fluorescent intensities that were at least 2.5 times above the background. Incorporation lengths were measured using the full-width-half-maximum (FWHM) of the intensity profiles; only incorporation stretches of at least 300 nm length were taken into account, based on the resolution limit of the microscope. For fracture experiments, images were taken every 5 seconds. Movies were processed to improve the signal-to-noise and signal-to-background ratios (smooth and subtract background functions of ImageJ, version 1.54j). Lengths of damaged regions were determined by line scans along the microtubules in the frame just prior to fracture; regions corresponding to a drop of at least 50% of the fluorescence intensity of the intact lattice were considered damaged.

### Quantification Of the Number of Incorporated Dimers in Incorporation Stretches

To estimate the number of incorporated dimers, we took the images from our incorporation experiments where we occasionally observed microtubule growth beyond the cap with 100% red-labeled tubulin that was used during the tubulin incorporation as reference stretch (Fig. 3d). We then used the integrated fluorescence of the reference stretch (original lattice) to estimate the number of fluorescent dimers in the incorporation site (normalized lattice fluorescence) (Fig. 3e). We performed this analysis for ten frames per incorporation stretch analyzed and averaged the values.

### Single-Molecule Photobleaching Experiment and Analysis

Tau proteins were diluted in cold 1x BRB80 to a concentration of a few tenths of micromolar and centrifuged to remove aggregates (10 min/4°C /215,000 x g in Type 70 Ti rotor [Beckman]). Prior to use, Tau proteins were further serial diluted to a final concentration of 50 pM. In the flow chamber, 50 μl of the diluted 2N4R-GFP Tau was perfused for 5 minutes at room temperature. The chamber was then washed three times with 100 μl 1x BRB80 to remove unbound proteins and then sealed with Valap. Images were recorded in continuous streaming mode for 2 minutes with 300ms exposure time and high laser power to photobleach the isolated molecules.

For analysis, a region of interest (ROI) with an uniform illumination was selected and the stack was cropped. The fluorescence intensity fluctuations over time of each fluorescent Tau molecule were obtained using Stowers Institute ImageJ Plugin (Fig. 1a, SI).

### Single-Molecule TIRF Microscopy

An objective-based orbital TIRF microscope with single-fluorophore sensitivity was used for the detection of fluorescently labeled microtubules and Tau molecules. Seeds were elongated with a mix containing 11 μM of tubulin (20% labeled with ATTO-565) in 0.7x BRB80 and 0.38x MAP Buffer (500mM Phosphate Buffer, 1M KCl, 10mM DTT, pH 7.9) supplemented with 1 mM GTP, an oxygen scavenger cocktail (22 mM DTT, 1.2 mg ml^−1^ glucose, 8 μg ml^−1^ catalase and 40 μg ml^−1^ glucose oxidase), 1 mg/ml casein and 0.033% (w/v) methyl cellulose (1,500 cP, Sigma) at 37°C. GMPcPP caps were grown by substituting in the before-mentioned buffer GTP with 0.5 mM GMPcPP (Jena Bioscience) and using 3 μM tubulin (100% labeled with ATTO-565) at 37°C. The assay buffer was supplemented with 50pM 2N4R-GFP Tau protein and free tubulin along with the same oxygen scavenger system as mentioned above to minimize bleaching of fluorophores. Additionally, a time-lapse video for 100 frames was recorded using the same laser power and same exposure obtained during the single-molecule photobleaching experiment.

### Statistical methods

All statistical analyses were performed using GraphPad Prism 9.5. The confocal microscopy-observed microtubule incorporation patterns were examined by concatenating all microtubules in random order and determining the center-to-center distance between two adjacent incorporation locations. To test the significance between incorporation lengths and distances between incorporation distributions obtained for the various conditions as well as for the lengths of the damaged regions (Fig. 2g) and number of incorporated protofilaments (Fig. 3f), we used the Mann-Whitney test (two-tailed) as a non-parametric alternative to a *t*-test, given that the distributions are non-Gaussian. To analyze the differences in the normalized lattice fluorescence (Fig. 3e), we used an unpaired *t*-test, given that the distributions appear Gaussian and with similar variances. Microtubule lifetime was quantified by manually counting the microtubule number at each frame within several fields of view, for 50 min. We used the Mantel-Cox log rank test to assess differences in microtubule survival curves.

### Coarse-Grained (CG) Molecular Dynamics Simulations

To estimate the free energy associated with tubulin dissociation in the presence and absence of tau, we performed CG molecular dynamics simulations. The atomistic starting structure of tau, spanning three tubulin monomers, was obtained from the protein data bank (PDB ID: 6CVN^24^).

The atomistic structure was converted to a CG representation using the Martinize script^47^ and modeled with the Martini-3 force field^48^. The structure of the monomers was preserved using an elastic network between backbone beads (force constant: 700 kJ mol^-1^ nm^-2^). The CG tau-tubulin complex was placed in a simulation box with dimensions of 20 nm x 20 nm x 20 nm, such that the microtubule would be oriented along the z-axis. The box was solvated with 62,508 standard Martini water beads and neutralized by adding 50 Na^+^ ions. The system was energy minimized using the steepest-descent algorithm. During equilibration, Lennard-Jones and Coulomb interactions were treated with a cutoff of 1.1 nm. The temperature was maintained at 310 K using velocity rescaling^49^ with a coupling constant of 0.1 ps, while the pressure was kept at 1 bar using the isotropic Berendsen barostat^50^ with a coupling constant of 6 ps. Bonds were constrained using the LINCS algorithm^51^. Equilibration was performed for 20 ns with the GROMACS 2021.1 simulation package^52^, with position restraints applied only on tau.

The final structure from the equilibration was used to generate starting configurations for umbrella sampling (US) simulations. The reaction coordinate was defined as the center-of-mass distance between adjacent monomers. A monomer was pulled away by 3 nm over 200 ns using an umbrella potential, where pull forces were applied only along the z-direction. A pull rate of 1.5 x 10^-5^ nm ps^-1^ and a force constant of 100000 kJ mol^-1^ nm^-2^ were applied. During pulling and US simulations, we applied position restraint potentials to avoid rotations of the overall monomer and to avoid that the monomers would drift in the x-y plane, because such motions could lead to sampling problems. Accordingly, except for the interface residues of the adjacent tubulin monomers which were fully flexible (residues 130–133, 163, 199, 241–265, 321–337, 345–355 of monomer 2 and residues 70–78, 95–102, 175–183, 206–224, 393–413 of monomer 3) position restraints were applied in x- and y-direction on backbone beads of all other residues with a force constant of 500 kJ mol^-1^ nm^-2^. No restraints were applied to the tau residues interacting with the pulled monomer 3, while the rest of tau residues had backbone position restraints with a force constant of 500 kJ mol^-1^ nm^-2^, thereby restraining the interfaces of tau with monomers 1 and 2. All other simulation parameters were identical with those used during equilibration.

From the pulling simulations, 170 structures were extracted for US simulations. The US windows were spaced at 0.015 nm with a force constant of 10000 kJ mol^-1^ nm^-2^ up to a distance of approximately 1.05 nm between the adjacent monomers. At larger distances, the spacing was set to 0.02 nm with a force constant of 5000 kJ mol^-1^ nm^-2^. Each US window was simulated for 100 ns using GROMACS 2021.1^52^ with a time step of 20 fs. The potential of mean force (PMF) was calculated after skipping the first 40 ns of each window for equilibration with the Weighted Histogram Analysis Method implementation gmx wham of GROMACS 2021.1^53,54^. Statistical errors were estimated using 50 rounds of bootstrapping of complete histograms. The PMF for the system without tau followed the same protocol, except that tau was removed from the equilibrated structure.

## Author contributions

Contributing authors (alphabetic order):

Biswas, Subham

Diez, Stefan

Grover, Rahul

Grünewald, Mona

Hub, Jochen S.

<u>John, Karin</u>

Poojari, Chetan S.

Reuther, Cordula

Shaebani, M. Reza

<u>Schaedel, Laura</u>

Zablotsky, Amir

LS, KJ and SD conceived and guided the project. LS, SD, SB, MG, RG and CR designed the experiments. SB carried out the experiments. MG purified tubulin. RG and CR provided tau and related expertise. AZ and KJ designed and carried out the kinetic Monte Carlo simulations. JSH and CSP designed and carried out the molecular dynamics simulations. MRS analyzed the single molecule experiments. LS, SD and KJ wrote the manuscript. All authors provided critical feedback and helped to shape the research and analysis.

## Acknowledgements

SB and LS were supported by the Deutsche Forschungsgemeinschaft (DFG, German Research Foundation; grant SFB 1027/A13) and through Saarland University’s NanoBioMed Young Investigator Grant awarded to LS. RG, CR and SD were supported by the DFG grant SFB 1027/A8. CSP and JSH were supported by the DFG grants SFB 1027/B7 and INST 256/539-1. MRS acknowledges support through the DFG grant SFB 1027/A7. KJ and AZ were supported through the Idex Université Grenoble-Alpes. The numerical computations by AZ and KJ were performed using the Cactus cluster of the CIMENT infrastructure, which was supported by the Rhône-Alpes region. The authors are grateful to Philippe Beys, who manages the cluster.

## Ethics declarations

The authors declare no competing interests.

