## Supplementary Information for "Tau accelerates tubulin exchange in the microtubule lattice"

**Affiliations**:

### Supplementary Information

#### Kinetic Monte Carlo (kMC) Model

To study the dynamics of microtubules containing $V_1$ and $V_2$ defects we designed a kinetic Monte Carlo (kMC) model on the scale of the tubulin monomer (Fig. 4b). With this model we explored the following hypothesis. Monomer vacancies are potentially weak spots in the microtubule lattice, since surrounding dimers are missing neighbors. Furthermore, the shift in the A-lattice seam across one protofilament (type $V_1$ vacancy) and the introduction of two new A-lattice seams (type $V_2$ vacancy) lead to a stress localization, which renders dimers in the vicinity of the defect less stable compared to the perfect lattice[1,2]. The tubulin incorporation visible in the experiments results from the mobility of monomer vacancies as depicted in Fig. 4e. Due to the lattice anisotropy, vacancies will preferentially move along a protofilament. If A-lattice contacts are less stable than B-lattice contacts (of the order of 1 kT[3–5]), $V_1$ and $V_2$ vacancies will behave differently: While $V_1$ defects will exhibit a mainly diffusive behavior, $V_2$ defects will exhibit a mainly ballistic motion in the direction of the additional A-lattice seam. Using kMC simulations we investigated the dynamics of $V_1$ and $V_2$ defects depending on the microtubule lattice properties (binding energy and anisotropy, localized lattice destabilization at the defect).

#### Model Description

kMC simulations were performed using a rejection-free random-selection method[6] and using custom written codes in C (Apple clang version 11.0.3). Statistical analysis was performed using C or R (R 4.0.0

for Mac OS X, https://www.R-project.org/). All model parameters are shown in Table S1 and Table S2 or are stated in the legends.

*Lattice structure:*

We model the canonical microtubule lattice (13 protofilaments, three-start left-handed helix[7,8]) as a square lattice on the scale of the tubulin monomer, i.e. each dimer has two longitudinal and 4 lateral (monomer) neighbors. The lattice is periodic in a direction perpendicular to the long axis of the microtubule with an offset of 3 monomer lattice sites to reproduce the seam structure (Fig. 4b). Lattice sites can be either empty or occupied by tubulin monomers. Lattice occupation occurs only via dimers, either bound to GTP (T) or GDP (D). Dimers interact with other dimers on nearest-neighbor lattice sites via attractive interactions, characterized by bond energies $\Delta G_1$ and $\Delta G_2$ for longitudinal and lateral bonds, respectively. $\Delta G_{tot}=2(\Delta G_1+\Delta G_2)$ denotes the total bond energy. Longitudinal T-T bonds are further stabilized by the energy $\Delta G^T_1$ [9–11]. We assume a lattice anisotropy in the binding energies of $A=\Delta G_1/\Delta G_2>1$ that is, longitudinal contacts are more stable than lateral contacts in the GDP lattice, consistent with estimates obtained by kinetic modeling of the microtubule tip dynamics[12] and molecular dynamics simulations[13]. The lattice may contain monomer sized vacant sites (Fig. 4b), which are dynamic (Fig. 4e). If such a monomer vacancy is adjacent to an A-lattice seam, the seam is shifted by a protofilament to the left or right, while the number of A-lattice contacts does not change in the lattice. If a monomer vacancy is placed amid the B-lattice, two new seams will originate from this vacancy, either in the direction towards the microtubule plus-end or the microtubule minus-end. For type 2 vacancies we distinguish between vacancies creating two new seams towards the plus end ($V_{2+}$) and towards the minus-end ($V_{2-}$).

*Kinetic Transitions:*

We consider the following transitions (Fig. 4b). Free GTP dimers can attach on vacant lattice dimer sites with rate constant $k_{on}$, if at least one neighboring lattice site is already occupied by a dimer. We assume an infinite reservoir of free GTP-tubulin dimers. We do not consider the attachment of GDP-dimers. Lattice bound GTP dimers may hydrolyze into GDP dimers with rate constant $k_{hy}$ if they form a longitudinal contact with another GTP or GDP dimer towards the microtubule plus-end[3]. Bound GTP and GDP dimers can detach from the lattice with rate constant $k_{off}$. depending on their lattice environment

$$k_{off} = \frac{1}{\tau} e^{\beta(\Delta G_b - \Delta G*)} (1)$$

where $\beta=1/(kT)$ is the inverse of thermal energy, $\Delta G_b$ denotes the binding energy and $\Delta G^*$ denotes a reference binding energy of a GDP-dimer associated with the off rate constant $k_{off}=1/\tau$.

*Handling of A-lattice contacts:*

At A-lattice contacts we count the number of monomer neighbors in the lateral direction. Each monomer contact counts with a binding energy of $(\Delta G_2+\Delta G_S)/2$, where $\Delta G_S$ denotes a weak destabilization of A-lattice contacts w.r.t. B-lattice contacts.

*Stress accumulation at monomer defects:*

Longitudinal transitions from A- to B-lattice contacts associated with a monomer vacancy are penalized by a weak destabilizing contribution $\Delta G_v>0$ (Fig. 4b and Fig. S4). Dimers in contact with several monomer vacancies are destabilized by summing up over all contributions. If a dimer in contact with a monomer vacancy leaves the lattice, the defect stress is released and the remaining dimers in contact with the monomer vacancy are stabilized.

*Effect of tau:*

In our experiments we consider sufficiently low concentrations of tau, where tau does not form domains, i.e. the fluorescence of tau is homogeneously distributed along the microtubule. In this case tau is rapidly diffusing (~0.2…0.3 $\mu m^2/s$ corresponding to a hopping rate of ~3000-5000 $s^{-1}$ between lattice sites) with a short residence time (~0.04...4 s corresponding to an off-rate constant of 0.25-to 25 $s^{-1}$) on microtubules (see Fig. S1 and Siahaan et al., 2019; Janning et al., 2014[14,15]). Compared to the tau dynamics, the tubulin dimer dynamics in the microtubule lattice is slow. For an incorporation length of 1 µm per 15 min via a ballistic defect motion we can roughly estimate a hopping rate constant for the defect of 0.1 $s^{-1}$ (assuming a dimer length of 8 nm), which is slower than any other rate constant associated with the tau dynamics. Therefore, we included the effect of tau on the microtubule lattice via an effective approach, where we assume that tau globally modifies lattice parameters (total binding energy $\Delta G_{tot}$, lattice anisotropy A), without modeling directly the tau dynamics. This approach has been successfully applied to model the destabilizing effect of molecular motors on the lattice[16]. For simplicity we assume that tau affects the GTP- and GDP-lattice in the same way, i.e. we ignore here deliberately the work by Castle and colleagues[17] where it was shown that tau interacts with the GTP-lattice more weakly than with the GDP-lattice.

*Lattice setup of simulations with stochastically placed monomer vacancies:*

For simulations of microtubule fracture and free tubulin incorporation we used a microtubule lattice with a stochastic placement of monomer vacancies with a given frequency. In experiments tubulin incorporations were observed with a spatial frequency of ~0.1 $\mu m^{-1}$. In our microtubule setup we therefore assumed that $V_1$ vacancies are placed with a spatial frequency of 0.05 $\mu m^{-1}$ at an existing seem and $V_{2+}$ vacancies are placed at a spatial frequency of 0.05 $\mu m^{-1}$ in the B-lattice. Thereby we constructed the microtubule from the minus-end towards the plus-end, i.e. the occurrence of a $V_1$ vacancy shifts the seam by one protofilament in the direction of the microtubule plus-end. The occurrence of $V_{2+}$ defects creates two new seams in the B-lattice which extend towards the microtubule plus-end, and which can serve as new locations for $V_1$ defects. Two A-lattice seams which are separated by a single protofilament may vanish due to the occurrence of a $V_{2-}$ defect with a spatial frequency of 0.5 $\mu m^{-1}$. The choice of the spatial frequency for the occurrence of $V_{2-}$ defects is somewhat arbitrary and is designed to create stretches of multi-seam structures which are shorter than the microtubule length of ~10 $\mu m$.

Applying this procedure to 1000 microtubule of length 10 $\mu m$ results in the following spatial frequencies: 0.0601 $\mu m^{-1}$ ($V_1$), 0.0476 $\mu m^{-1}$ ($V_{2+}$), 0.0374 $\mu m^{-1}$ ($V_{2-}$) and a total frequency of defects of 0.1451 $\mu m^{-1}$. All simulations using microtubules with stochastic placement of monomer vacancies used exactly the same microtubule ensemble unless stated otherwise.

*Convolution of the positions of incorporated dimers to mimic the effect of the optical point spread function:*

To construct incorporation kymographs and to determine the size of the "visible" incorporations (Fig. 4 f, g in the main text and Fig. S8 and Fig. S9) we convoluted the longitudinal positions of incorporated dimers with a point spread function (FWHM=150 nm) and added over the contributions of all incorporated dimers. Only those incorporations were counted as visible which exceeded a threshold corresponding to ⅓ of the intensity of one fully replaced protofilament. Furthermore, as in the experiments, only those incorporations with a minimum length of 300 nm were counted, i.e. the simulated length distributions (Fig. 4g, right, and Fig. S8 and Fig. S9) only contain incorporation spots with a minimum length of 300 nm.

**Determining GDP-lattice parameters from microtubule fracture experiments**

In a first set of simulations, we investigated the fracture of microtubules in the absence of free tubulin depending on the lattice parameters (total binding energy of the GDP lattice $\Delta G_{tot}$ =2($\Delta G_1$+$\Delta G_2$) and lattice anisotropy A=$\Delta G_1/\Delta G_2$ for the GDP B-lattice, destabilization of lateral A-lattice contacts $\Delta G_S$). To that end we simulated the fracture of 10 µm long microtubules containing stochastically placed defects of type $V_1$, $V_{2+}$, $V_{2-}$ (with a total spatial frequency 0.15 $\mu m^{-1}$ as described above) and determined the longitudinal extension of the lattice damage at fracture. Here we considered a microtubule as broken, when the damaged region extended over all 13 protofilaments. The defect energy $\Delta G_V$ was neglected here since it has only a weak influence on the fracture (the defect associated stress is released as soon as the first dimer leaves the defect site).

Fig. S5a shows the size of damage at fracture depending on the lattice parameters ($\Delta G_{tot}$, A, $\Delta G_S$). From these simulations we identified a range of possible parameter combinations (Table S2) corresponding to the microtubule fracture experiments in the absence (control, damage size ~1 µm) and presence of tau (tau, damage size ~3 µm). The time scale was adjusted to match the experimentally measured time to fracture in the absence of tau (~15 min). Furthermore, we found that a weak stabilization of the total binding energy by -0.2 kT is sufficient to describe the increased time of microtubule survival in the presence of tau (~20 min). The microtubule survival curves for various lattice parameters are shown in Fig. S5b.

**Measuring tubulin incorporation at defect sites**

Without restricting generality we considered total GDP lattice energies in the range of -44…-36 kT, similar to values in the literature[12,16,18–22]. We then investigated the vacancy dynamics in the presence of free tubulin, i.e. tubulin dimers in the vicinity of the defect may leave the lattice and GTP tubulin dimers from the solution may attach to the lattice at vacant dimer sites as depicted in Fig. 4e. The additional parameters associated with the defect dynamics (GTP-dimer attachment rate constant $k_{on}$, longitudinal GTP-lattice stabilization $\Delta G_1^T$, hydrolysis rate constant $k_{hy}$) were adjusted such that the microtubule tip exhibits a dynamic instability with a growth rate of ≤ 1 µm/min close to the critical $k_{on}$

(Fig. S6 and Fig. S7). The on-rate constant for the investigation of the defect dynamics was chosen to be slightly smaller than the critical $k_{on}$-rate, as in the experiments.

In the absence of any lateral A-lattice seam destabilization ($\Delta G_S$=0 kT) both $V_1$ and $V_2$ vacancies incorporate free tubulin with the same dynamics (Fig. S8a, left). In general, defects incorporate more free tubulin in the presence of tau than in the absence due to increased lattice anisotropy in the presence of tau. In the absence of any defect stress ($\Delta G_V$=0 kT), incorporations are small < 100 nm. Upon destabilizing the dimers in the immediate vicinity of the defect ($\Delta G_V$>0 kT), tubulin is incorporated over longer stretches along a single protofilament. Simulating the incorporation of free tubulin into microtubules with stochastically placed defects leads, however, to a relatively broad spectrum of visible incorporations (Fig. S8a, right).

In the presence of a weak lateral A-lattice seam destabilization ($\Delta G_S$>0 kT) $V_2$ vacancies incorporate more tubulin than $V_1$ vacancies (Fig. S8 b, c) under control conditions and in the presence of tau, which also leads to a more complex distribution of visible incorporations into microtubules with stochastically placed defects. For example, for a defect destabilization of $\Delta G_V$=1.25 kT and an A-lattice seam destabilization of $\Delta G_S$=0.5 kT the simulated incorporation sizes are in the experimentally measured range. In this situation, in the absence of tau the majority of visible defects corresponds to $V_2$ defects, whereas $V_1$ defects are too small to be counted as incorporation. However, in the presence of tau, $V_2$ defects become longer and at the same time a larger portion of $V_1$ defects becomes visible. The incorporation behavior at V1 and V2 defects is robust and leads to comparable incorporations lengths for the tested lattice binding energies in the the range $\Delta G_{tot}$=-44…-36 kT, A-lattice destabilization $\Delta G_S$=0…1 kT and defect energy $\Delta G_V$=0…2 kT (see also Fig. S9).

**Table S1:**

| Parameter | Value | References/Remarks |
|---|---|---|
| total lattice binding energy $\Delta G_{tot}$ | -44 … -36 kT | Adapted in this model to match experimental data (microtubule fracture experiments). Comparable to Kononova et al., 2014; Ganser and Uchihashi, 2019[18,19]; and previous kMC models for the microtubule tip (vanBuren et al., 2002, Wu et al., 2009; Gardner et al.; 2011, Schaedel et al., 2019, Lecompte and John, 2023[12,16,20–22]) |
| lattice anisotropy A | 1.2 … 2.2 kT | |
| Seam penalty (lateral A-lattice contacts) $\Delta G_S$ | 0.0 … 1.0 kT | Adapted in this model to match experimental data. |
| Defect energy $\Delta G_V$ | 0.0 … 2.0 kT | |
| Longitudinal stabilization of T-T contacts $\Delta G_1^T$ | -7.2 … -3.6 kT | 10% … 20% of total binding energy |
| Stabilization of the total lattice binding energy $\Delta G_{tot}$ by tau | -0.2 kT | Adapted in this model to match experimental data (increase in time to microtubule fracture in the absence of free tubulin). |
| Hydrolysis rate constant | 0.25 … 1 $s^{-1}$ | Adapted in this model, comparable to Melki et al., 1996[23] and other kMC models (vanBuren et al., 2002, Schaedel et al., 2019, Lecompte and John, 2023[12,16,22]) |
| Effective tubulin on-rate for incorporation experiments | 1 … 10 $s^{-1}$ | Adapted in this model to be slightly below the critical on-rate for polymerization, comparable to vanBuren et al., 2002, Wu et al., 2009[12,20]) |
| tubulin off-rate constant (reference state) $k^*_{off}=1/\tau$ | 7 … 50 $s^{-1}$ | Adapted in this model to match experimental data (time to microtubule fracture in the absence of free tubulin and tau). |

*Table S1: Ranges of parameter values used in the kMC model if not stated otherwise.*

**Table S2:**

| Label | Lattice Parameters | Tubulin Incorporation Parameters |
|---|---|---|
| I: control | $\Delta G_{tot}$=-36.0 kT, $\Delta G^*$=-18.0 kT, $\Delta G_S$=0.0, A=1.55, τ=0.094 s | $\Delta G_1^T$=-3.6 kT, $k_{hy}$=1 $s^{-1}$, $k_{on}$=1 $s^{-1}$ |
| I: tau | $\Delta G_{tot}$=-36.2 kT, $\Delta G^*$=-18.0 kT, $\Delta G_S$=0.0, A=2.15, τ=0.094 s | |
| IIa: control | $\Delta G_{tot}$=-36.0 kT, $\Delta G^*$=-18.0 kT, $\Delta G_S$=0.5, A=1.50, τ=0.117 s | $\Delta G_1^T$=-3.6 kT, $k_{hy}$=1 $s^{-1}$, $k_{on}$=1 $s^{-1}$ |
| IIa: tau | $\Delta G_{tot}$=-36.2 kT, $\Delta G^*$=-18.0 kT, $\Delta G_S$=0.5, A=2.10, τ=0.117 s | |
| IIb: control | $\Delta G_{tot}$=-36.0 kT, $\Delta G^*$=-18.0 kT, $\Delta G_S$=0.5, A=1.50, τ=0.117 s | $\Delta G_1^T$=-7.2 kT, $k_{hy}$=0.25 $s^{-1}$, $k_{on}$=1 $s^{-1}$ |
| IIb: tau | $\Delta G_{tot}$=-36.2 kT, $\Delta G^*$=-18.0 kT, $\Delta G_S$=0.5, A=2.10, τ=0.117 s | |
| III: control | $\Delta G_{tot}$=-36.0 kT, $\Delta G^*$=-18.0 kT, $\Delta G_S$=1.0, A=1.40, τ=0.152 s | $\Delta G_1^T$=-3.6 kT, $k_{hy}$=1 $s^{-1}$, $k_{on}$=1 $s^{-1}$ |
| III: tau | $\Delta G_{tot}$=-36.2 kT, $\Delta G^*$=-18.0 kT, $\Delta G_S$=1.0, A=2.00, τ=0.152 s | |
| IV: control | $\Delta G_{tot}$=-40.0 kT, $\Delta G^*$=-20.0 kT, $\Delta G_S$=0.5, A=1.35, τ=0.047 s | $\Delta G_1^T$=-4.0 kT, $k_{hy}$=1 $s^{-1}$, $k_{on}$=6 $s^{-}$ |
| IV: tau | $\Delta G_{tot}$=-40.2 kT, $\Delta G^*$=-20.0 kT, $\Delta G_S$=0.5, A=1.85, τ=0.047 s | |
| V: control | $\Delta G_{tot}$=-44.0 kT, $\Delta G^*$=-22.0 kT, $\Delta G_S$=0.5, A=1.25, τ=0.019 s | $\Delta G_1^T$=-4.4 kT, $k_{hy}$=1 $s^{-1}$, $k_{on}$=10 $s^{-}$ |
| V: tau | $\Delta G_{tot}$=-44.2 kT, $\Delta G^*$=-22.0 kT, $\Delta G_S$=0.5, A=1.65, τ=0.019 s | |

*Table S2: Parameter combinations used in the kMC model if not stated otherwise.*

**Fig. S1**

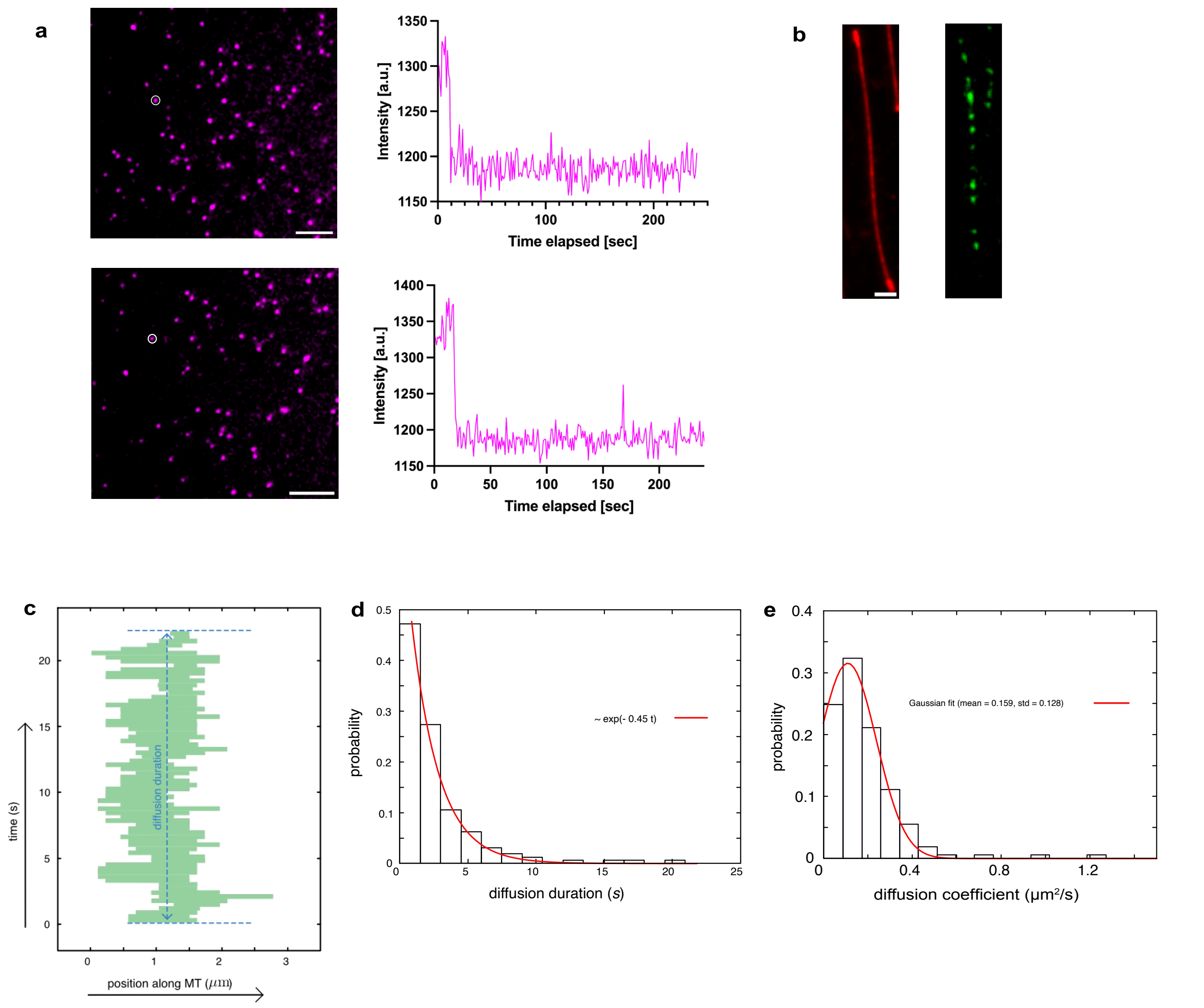


**Fig. S1:**

a. Example images for single-molecule photobleaching at 50pM 2N4R Tau-GFP exposed for 300ms along with corresponding fluctuations of fluorescence intensity of Tau-GFP over time. One-step bleaching indicates the presence of monomeric protein. Scale Bar: 5µm.
b. Example image of microtubule (red) and snapshot of 50pM fluorescently labeled tau (green) moving along the microtubule lattice. The mean fraction of tau-covered microtubule is 0.191 ± 0.008. Scale bar: 3 µm.
c. A sample kymograph of the movement of tau along a microtubule. The kymograph was reconstructed by varying the intensity threshold parameter.
d. Probability distribution of the diffusion duration. The mean duration of diffusion is 3.7 ± 0.7 s. The line represents an exponential fit.
e. Probability distribution of the diffusion coefficient. The mean diffusion coefficient corresponds to 0.21 ± 0.03 µm$^2$/s. The line represents a Gaussian fit.

Fig. S2

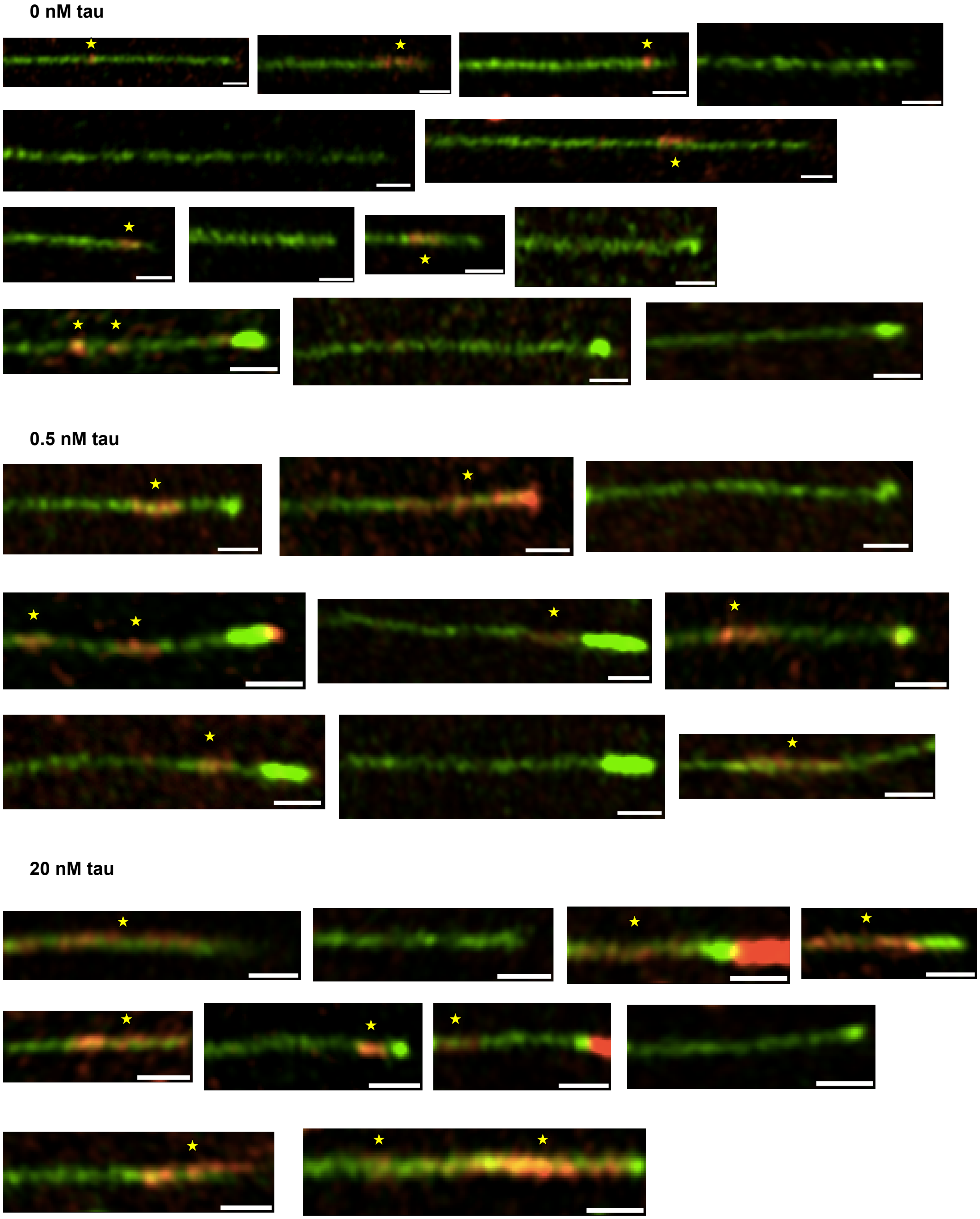


**Fig. S2:**
Example images of incorporation along the microtubule lattice in 15 mins at different tau concentrations. Scale bar: 1 μm.

**Fig. S3**

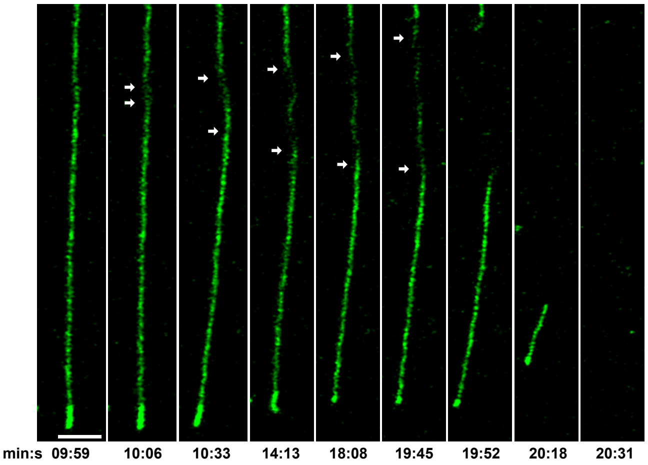


**Fig. S3:**
Example image sequence of the spreading of a damaged region in the absence of free tubulin (fracture experiment). Scale bar: 3 µm.

### Fig. S4

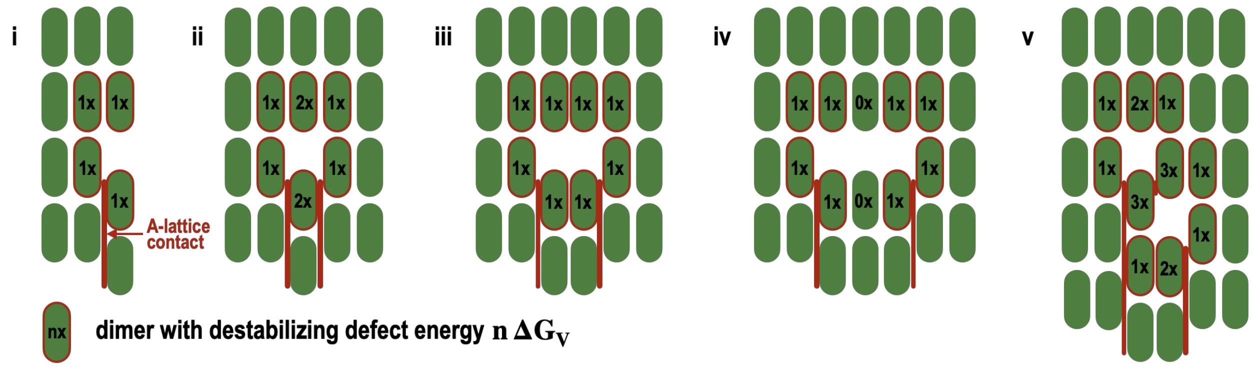


**Fig. S4:**

Schematic representation of the kMC model setup. Shown is the distribution of the defect energy in the neighborhood of monomer vacancies. i: Principal element of a monomer vacancy, where a vertical line of lateral A-lattice contacts (thick red line) transforms into a vertical line of lateral B-lattice contacts from bottom to top. All dimers, which are destabilized by vertical A<->B lattice transition are highlighted by a red boundary. ii-v: Examples of lattice configurations and the corresponding defect energies per dimer. Note that in the presence of several A<->B lattice transition the defect energy per dimer is obtained by summation over the contribution from each vertical A<->B transition.

**Fig. S5**

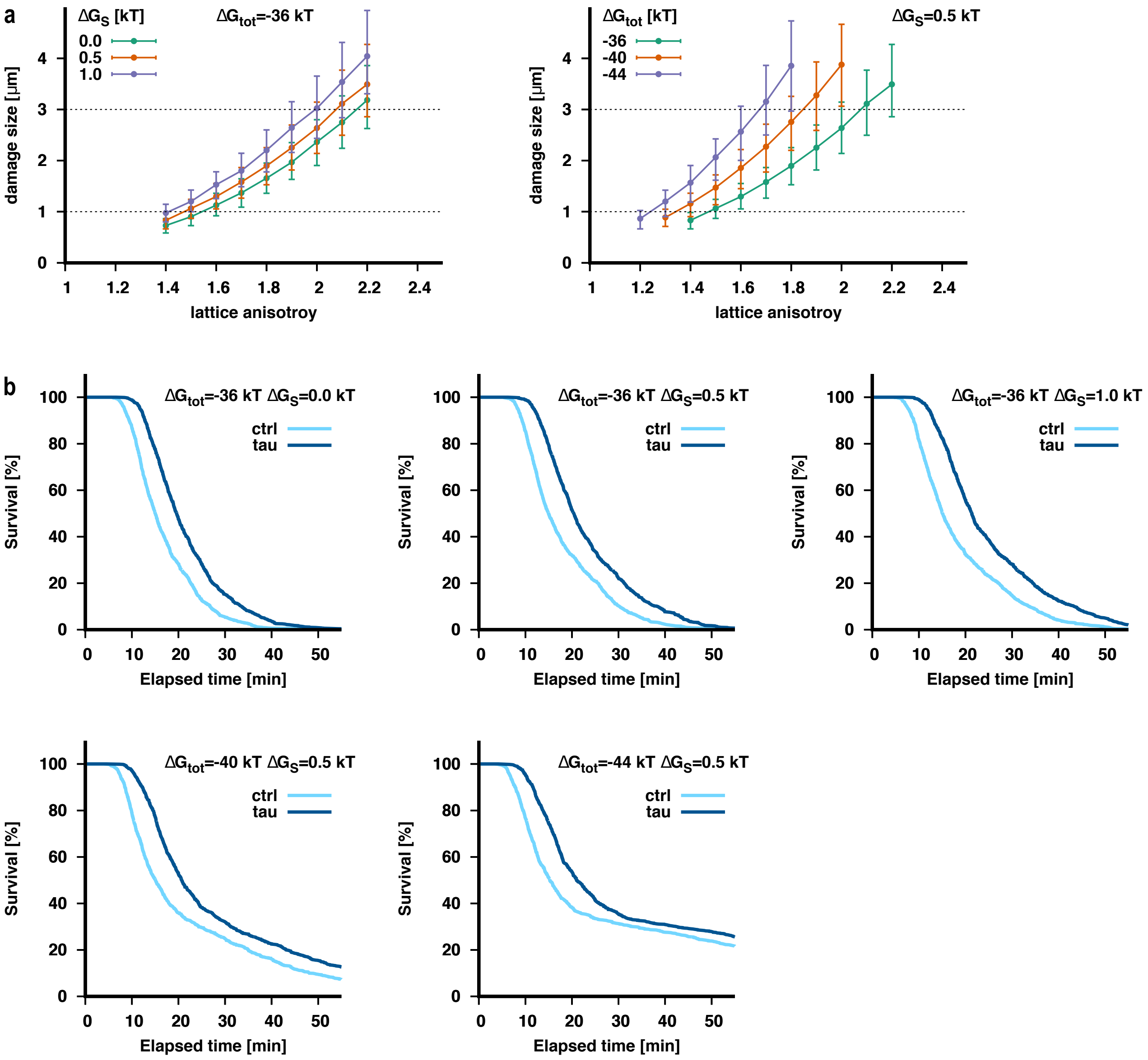


**Fig. S5: Simulation of microtubule fracture in the absence of free tubulin dimers.**

**a.** Longitudinal extension of damage at complete microtubule fracture depending on the GDP-lattice parameter (total binding energy $\Delta G_{tot}$, lattice anisotropy A, and lateral A-lattice destabilization $\Delta G_S$). Simulated microtubules are 10 $\mu$m long with stochastically placed defects as described in SI with a special frequency of 0.15 $\mu m^{-1}$.

**b.** Survival curves of microtubules for various lattice parameters (indicated in each plot and in Table S2) in the absence and presence of tau.

Fig. S6

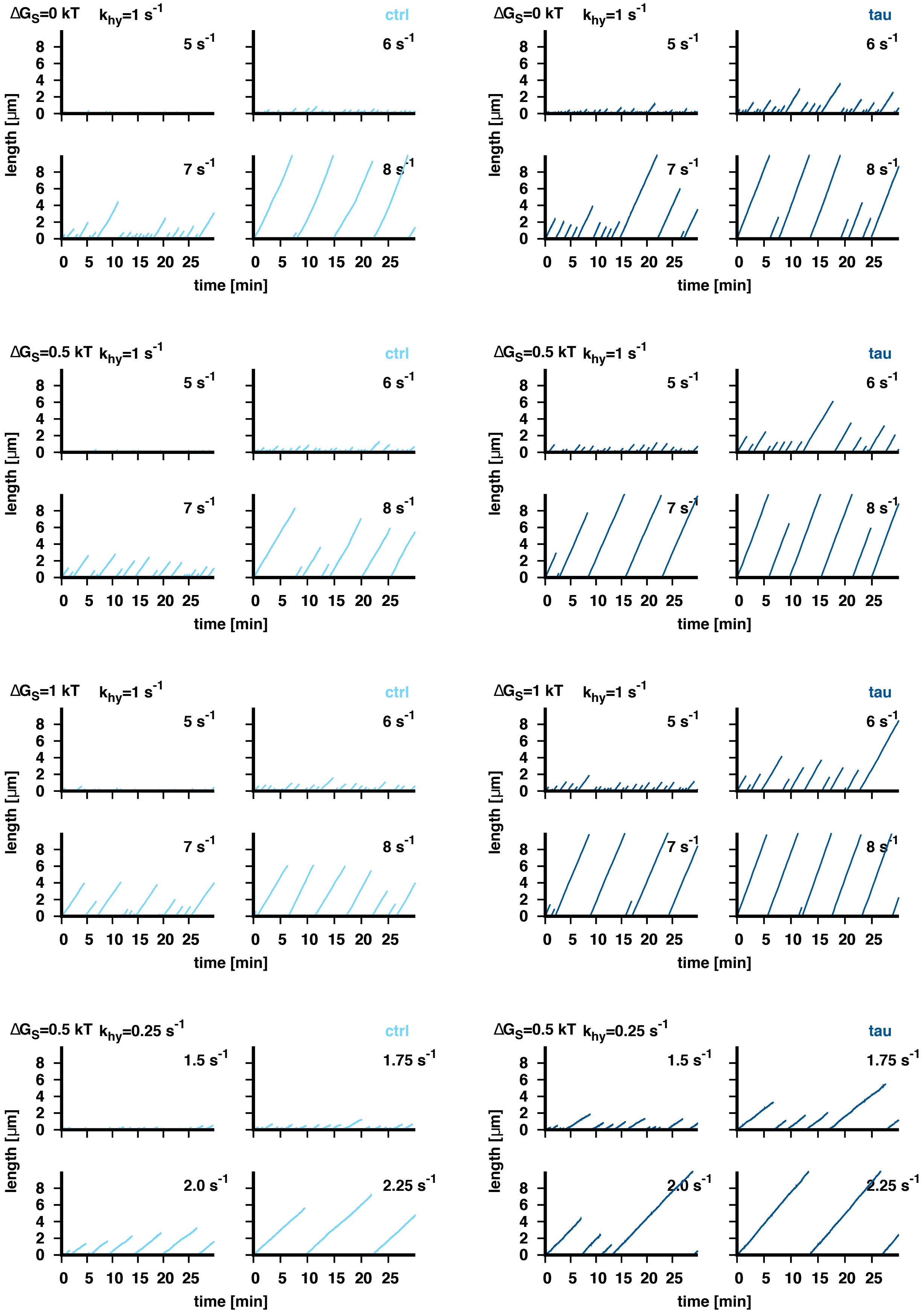
ΔGS=0 kT khy=1 s-1
ctrl
5 s-1
6 s-1
7 s-1
8 s-1
ΔGS=0 kT khy=1 s-1
tau
5 s-1
6 s-1
7 s-1
8 s-1
ΔGS=0.5 kT khy=1 s-1
ctrl
5 s-1
6 s-1
7 s-1
8 s-1
ΔGS=0.5 kT khy=1 s-1
tau
5 s-1
6 s-1
7 s-1
8 s-1
ΔGS=1 kT khy=1 s-1
ctrl
5 s-1
6 s-1
7 s-1
8 s-1
ΔGS=1 kT khy=1 s-1
tau
5 s-1
6 s-1
7 s-1
8 s-1
ΔGS=0.5 kT khy=0.25 s-1
ctrl
1.5 s-1
1.75 s-1
2.0 s-1
2.25 s-1
ΔGS=0.5 kT khy=0.25 s-1
tau
1.5 s-1
1.75 s-1
2.0 s-1
2.25 s-1
length [µm]
time [min]
0
2
4
6
8
0 5 10 15 20 25

**Fig. S6: Simulation of microtubule tip dynamics in the presence of free tubulin dimers.**

Shown are only growth phases. Catastrophe events are defined to occur when the microtubule extremity is exclusively composed of GDP-dimers. The subsequent phase of microtubule shortening with the formation of outcurling protofilaments is not simulated for simplicity. The total lattice binding energy is $\Delta G_{tot}$=-36 kT (control) and $\Delta G_{tot}$=-36.2 kT (tau). The remaining parameters are indicated in the top region of each plot and as summarized in Table S2.

### Fig. S7

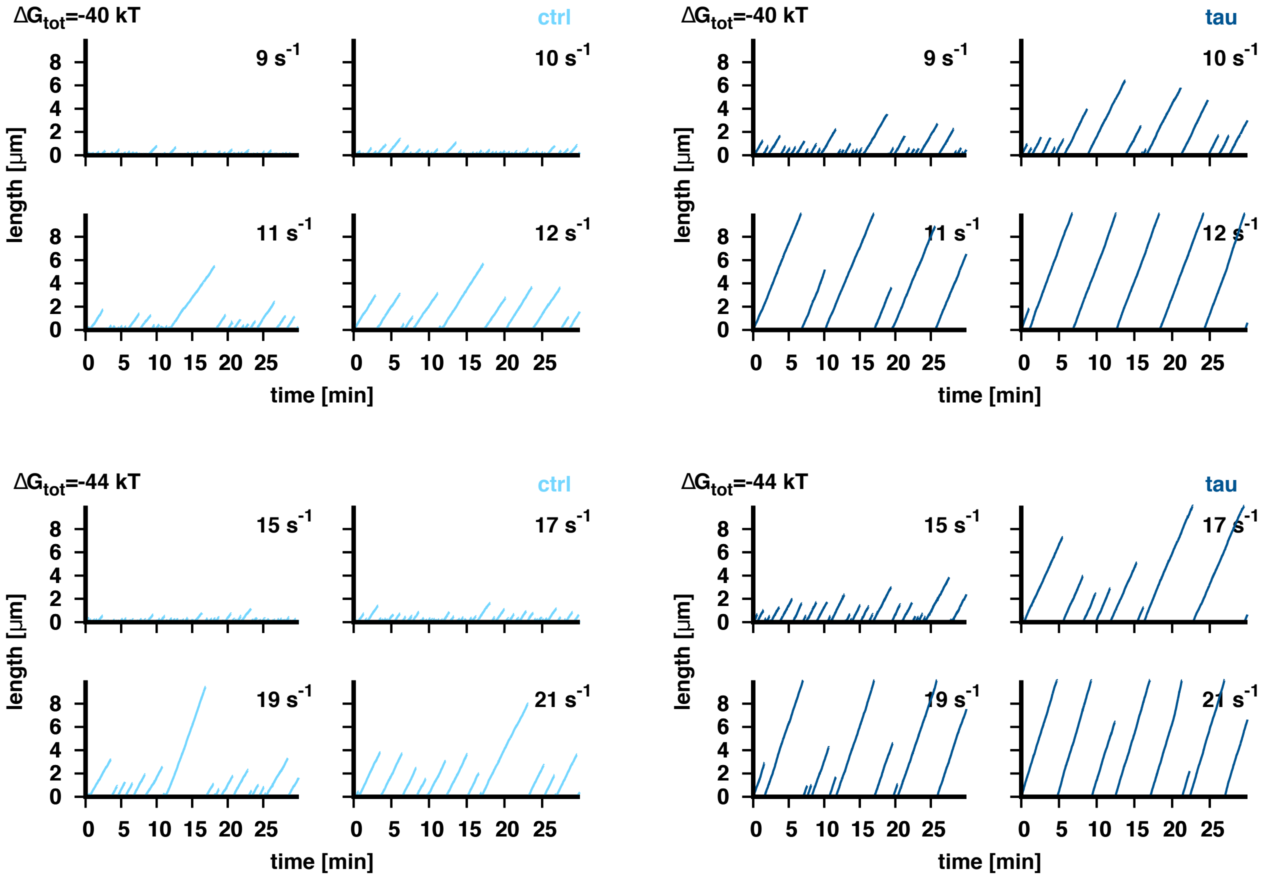


**Fig. S7: Simulation of microtubule tip dynamics in the presence of free tubulin dimers.**
Shown are only growth phases. Catastrophe events are defined to occur when the microtubule extremity is exclusively composed of GDP-dimers. The subsequent phase of microtubule shortening with the formation of outcurling protofilaments is not simulated for simplicity. The lateral A-lattice destabilizing energy is $\Delta G_S$=0.5 kT and the hydrolysis rate constant is $k_{hy}$=1 $s^{-1}$. The remaining parameters are indicated in the top region of each plot and as summarized in Table S2.

Fig. S8

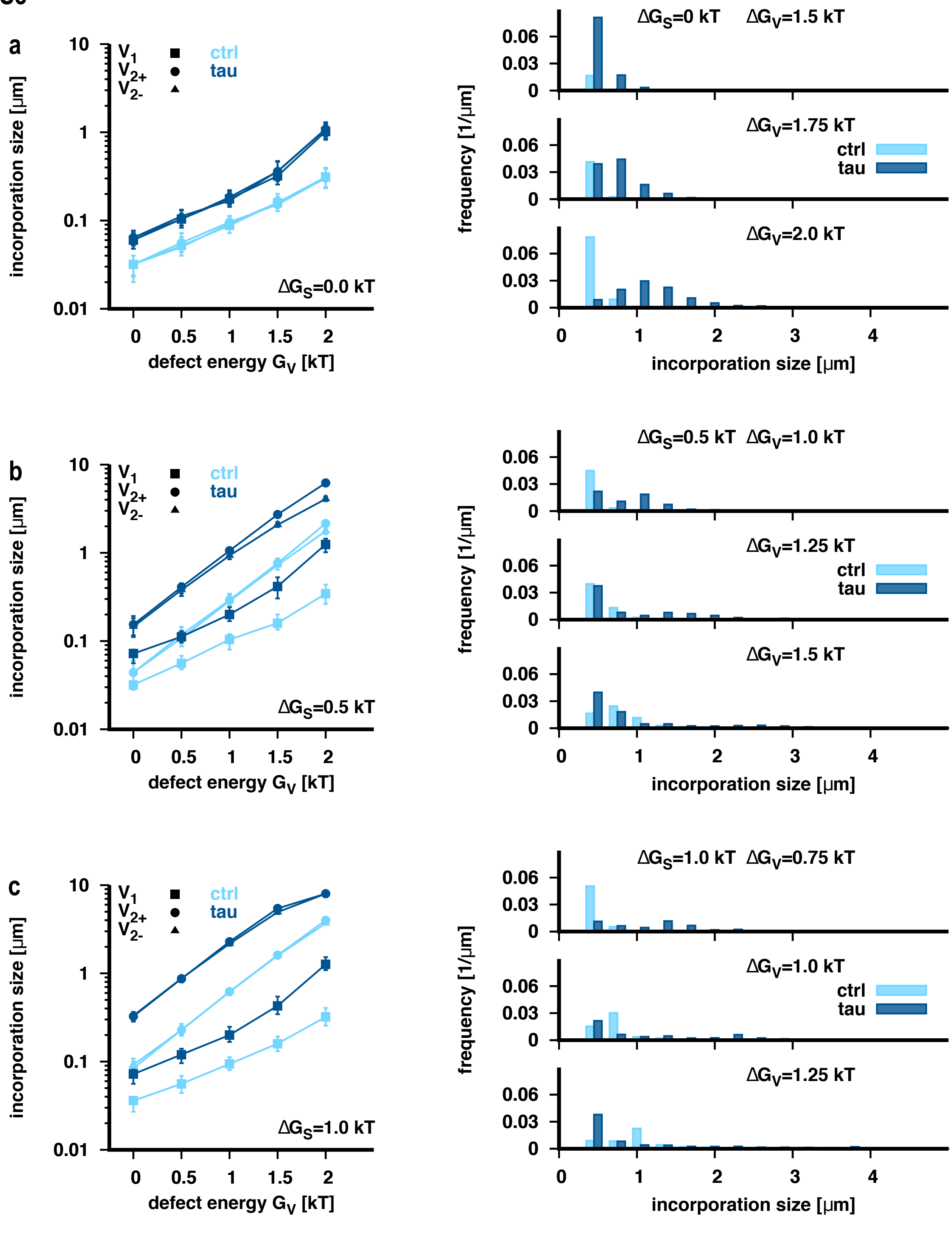


**Fig. S8: Simulation of free tubulin incorporation at defects sites.**
Shown are the "true" incorporation lengths for each defect type (left) depending on the defect energy $\Delta G_V$ and the size of "visible" incorporations (convoluted by the optical point spread function, right) for incorporation into microtubules with stochastically placed defects with defect frequency of 0.15 $\mu m^{-1}$. The total binding energy is $\Delta G_{tot}$=-36 kT. The remaining parameters are as indicated in each plot and in Table S2.

Fig. S9

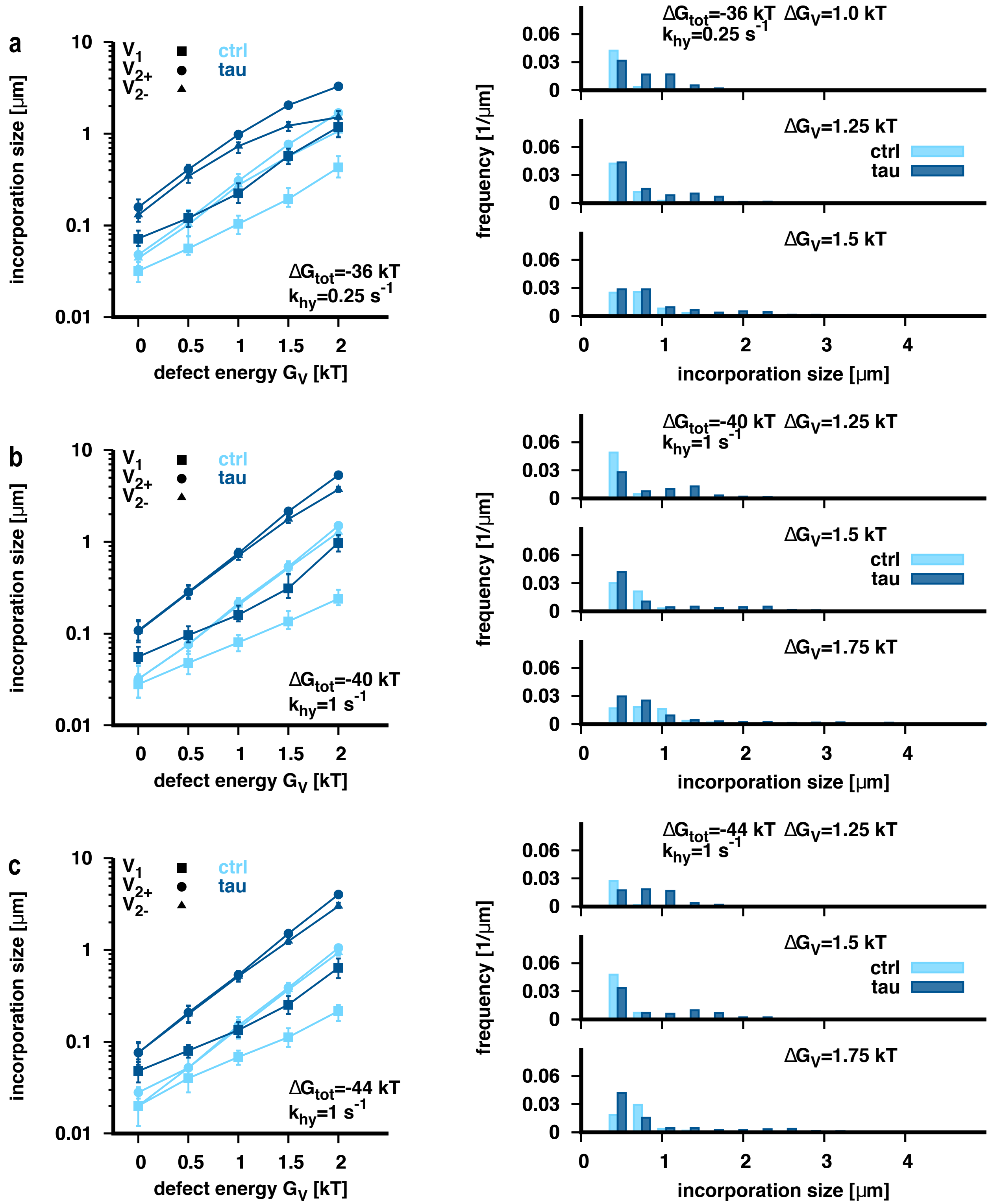


**Fig. S9: Simulation of free tubulin incorporation at defects sites.**
Shown are the "true" incorporation lengths for each defect type (left) depending on the defect energy $\Delta G_V$ and the size of "visible" incorporations (convoluted by the optical point spread function, right) for incorporation into microtubules with stochastically placed defects with defect frequency of 0.15 $\mu m^{-1}$. The lateral A-lattice seam destabilization is $\Delta G_S$=0.5 kT. The remaining parameters are as indicated in each plot and in Table S2.

**Fig. S10**

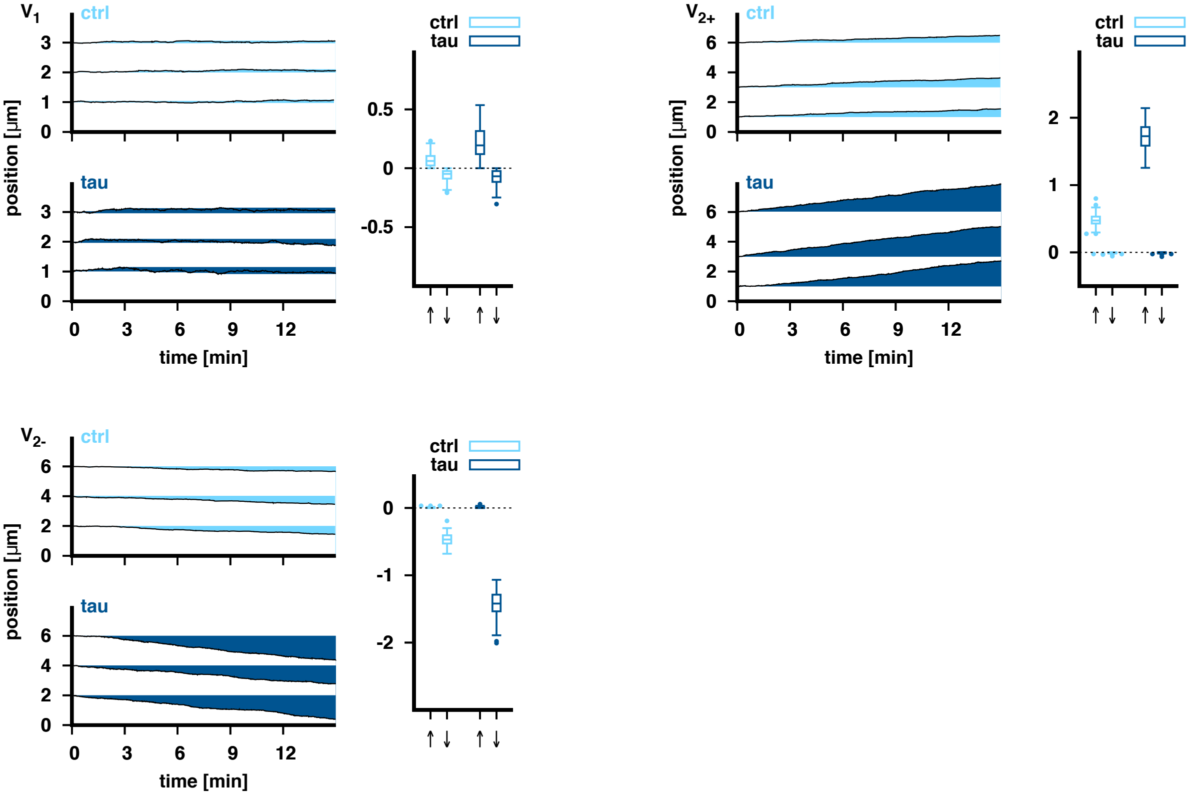


**Fig. S10:**

Example trajectories (left panels) for $V_1$, $V_{2+}$, $V_{2-}$ defects (as indicated in the left top corner of each plot) in the absence (ctrl, light blue color) and in the presence of tau (tau, dark blue color). The right panel shows the maximum and minimum positions where free tubulin was incorporated. The position of the defect at time zero is at zero. Shown are the median and the 25% and 75% quantiles. An asymmetry of maximum and minimum positions w.r.t. zero indicates that the defect motion has a ballistic component. Parameters are $\Delta G_S$=0.5 kT, and $\Delta G_{tot}$=-36 kT, A=1.5 (control) and $\Delta G_{tot}$=-36.2 kT, A=2.1 (tau) with $\Delta G_V$=1.25 kT and as indicated in Table S2.

<u>**References**</u>


1. Kovács, István and Hussein Amin El Sayed. "Point defects in metals." *Journal of Materials Science* 11, 529-559 (1976).
2. Pertsinidis, A. & Ling, X. S. Diffusion of point defects in two-dimensional colloidal crystals. *Nature* **413**, 147–150 (2001).
3. Nogales, E. A structural view of microtubule dynamics. *Cellular and Molecular Life Sciences (CMLS)* **56**, 133–142 (1999).
4. Katsuki, M., Drummond, D. R. & Cross, R. A. Ectopic A-lattice seams destabilize microtubules. *Nat Commun* **5**, 3094 (2014).
5. Harris, B. J., Ross, J. L. & Hawkins, T. L. Microtubule seams are not mechanically weak defects. *Phys. Rev. E* **97**, 062408 (2018).
6. Lukkien, J. J., Segers, J. P. L., Hilbers, P. A. J., Gelten, R. J. & Jansen, A. P. J. Efficient Monte Carlo methods for the simulation of catalytic surface reactions. *Phys. Rev. E* **58**, 2598–2610 (1998).
7. Mandelkow, E. M., Schultheiss, R., Rapp, R., Müller, M. & Mandelkow, E. On the surface lattice of microtubules: helix starts, protofilament number, seam, and handedness. *Journal of Cell Biology* **102**, 1067–1073 (1986).
8. Chrétien, D. & Wade, R. H. New data on the microtubule surface lattice. *Biology of the Cell* **71**, 161–174 (1991).
9. Alushin, G. M. *et al.* High-Resolution Microtubule Structures Reveal the Structural Transitions in αβ-Tubulin upon GTP Hydrolysis. *Cell* **157**, 1117–1129 (2014).
10. Molodtsov, M. I. *et al.* A Molecular-Mechanical Model of the Microtubule. *Biophysical Journal* **88**, 3167–3179 (2005).
11. Igaev, M. & Grubmüller, H. Microtubule instability driven by longitudinal and lateral strain

propagation. *PLoS Comput Biol* **16**, e1008132 (2020).
